# A simple platform for *in situ* DNA-PAINT optimization enables improved multiplexed super-resolution imaging in Gram-negative bacteria

**DOI:** 10.64898/2026.10.11.758320

**Authors:** Kilian Andress, Paul Said Köhler, Ruilan Xu, Alica Schmidt-Heydt, Andreas Diepold, Christoph Spahn

**Affiliations:** Rudolf Virchow Center for Integrative and Translational Bioimaging, University Würzburg, 97080 Würzburg, Germany; Institute of Applied Biosciences, Department Applied Biology, Karlsruhe Institute of Technology, 76131 Karlsruhe, Germany

## Abstract

DNA-PAINT enables super-resolution imaging with high multiplexing capacity, yet it is rarely used in nanoscale bacteriology. A likely reason is the lack of straightforward protocols and of systematic characterization of imager-docking strand pairs inside intact bacterial cells, whose envelope and dense cytoplasm differ markedly from both the DNA origami *in vitro* systems used to develop them, and eukaryotic cells. In this work, we establish a screening platform based on RNA-FISH, in which we extend the 16S rRNA-directed EUB338 probe with docking sequences of choice. This allows imager-docking pairs to be benchmarked directly inside *Escherichia coli* cells and characterized for parameters such as specificity, binding kinetics, and localization precision. We screened 12 different DNA-PAINT sequences and used an optimized set to perform four-target 3D Exchange-PAINT in single *E. coli* cells. We further applied our strategy to image three effector mRNAs in the Gram-negative pathogen *Yersinia enterocolitica* and quantified RNA clusters and their 3D distribution. Our work provides a framework and practical guidelines for high-quality 3D DNA-PAINT imaging in Gram-negative bacteria and should facilitate its broader adoption in nanoscale bacteriology.

## Introduction

Many of the processes that define bacterial physiology, such as chromosome organization, protein expression, secretion or cell wall synthesis, occur on length scales of tens of nanometers, well below the diffraction limit of conventional light microscopy [1]. Renewed interest in bacterial cell biology, driven in part by the search for new antimicrobial targets and a mechanistic understanding of host–pathogen interactions, has created strong demand for super-resolution methods that work reliably inside bacterial cells. The small size and high molecular crowding within bacterial cells require high- or super-resolution approaches to dissect molecular mechanisms and target distributions [1,2].

Among the available approaches, STED [3], SIM [4], and single-molecule localization microscopy (SMLM) techniques, such as *d*STORM, PALM, and (DNA-)PAINT [5–10], DNA-PAINT is particularly attractive for bacterial imaging. As single-molecule signal in DNA-PAINT is generated by transient hybridization of fluorescently labelled imager strands to complementary docking strands, photobleaching of individual fluorophores does not limit acquisition, and the technique routinely reaches near-molecular resolution under TIRF or HILO illumination [8,11,12]. Most importantly, Exchange-PAINT decouples the number of targets from the number of spectrally distinct fluorophores: by sequentially introducing different imager strands of different sequences, multiplexing capacity is set by the number of orthogonal imager–docking pairs rather than by the available dye palette [13]. This makes DNA-PAINT well suited to dissect the molecular organization of a system as small and densely packed as a bacterial cell, where many targets must be mapped within a volume of only a few femtoliters.

A growing library of imager strands has been developed for DNA-PAINT, including the original P-sequences [13], the repetitive, kinetically optimized R-sequences [14,15], fluorogenic imager strands [16], and left-handed DNA variants designed to expand the orthogonal pool and to reduce unspecific binding to nucleic acids [17,18]. The orthogonality of these pairs was characterized in detail at the single-molecule level on DNA origami [14,19] and, being a property of the hybridization sequences themselves, is expected to be largely independent of the cellular environment. What *is* environment-dependent, and what determines whether an imager is usable inside a bacterium, are the parameters that govern actual imaging performance: sequence-specific binding with minimal off-target signal, high fluorophore brightness and hybridization kinetics that provide well-separated single-molecule events [20].

Binding kinetics are typically studied in highly controlled systems, mostly DNA origami, which are stabilized by specific buffer conditions [8,14,19]. However, the results might not be transferrable to biological systems, which require imaging conditions adjusted to the more complex environment. Furthermore, in contrast to exposed DNA-origami, accessibility might be strongly altered in biological systems. This is particularly true for bacteria, in which accessibility is challenged by the two membranes and peptidoglycan layer as size-selective entry barriers [21–23].

Despite its huge potential, DNA-PAINT imaging, therefore, has remained a niche application in bacteriology. Cramer *et al*. used 2-plex DNA-PAINT to image the interplay of FtsZ and DivIVA during *Bacillus subtilis* sporulation [24]. Ermoli *et al*. employed a DNA-PAINT–based FISH approach to map effector mRNA localization in *Y. enterocolitica* in correlation to the membrane and DNA as subcellular landmarks [25]. Each of these studies relied on a small, pre-selected set of imager-docking strand pairs adopted from mammalian work, without a comparative evaluation of imager performance in the bacterial environment. As a result, the community currently has no systematic basis for choosing, or rejecting, imager strands for bacterial DNA-PAINT imaging.

Here we close this gap by establishing a system for quantitative *in situ* screening of imager-docking strand pairs within intact bacterial cells. The assay builds on rRNA-FISH against the 16S rRNA using the widely employed eubacterial probe EUB338-I [26,27] (from here on referred to as EUB338). This sequence targets a conserved, accessible loop shared by most members of the domain *Eubacteria* (Figure 1A). Extending the FISH sequence with docking strands allows for cost-effective and straight-forward screening of any DNA-PAINT sequence for application to intracellular bacterial targets. The simple protocol and high commercial availability of DNA oligomers facilitate tuning of docking and imager strands, e.g. by altering the number of repeats in repetitive docking sequences (R strands) as well as varying imager strand lengths to optimize binding kinetics for the biological system or target molecule. We developed an automated analysis pipeline to extract relevant parameters, such as theoretical and experimental localization precisions, ON-times and specificity, which enabled us to quantitatively assess the performance of docking-imager strand pairs under various conditions.

**Figure 1:**
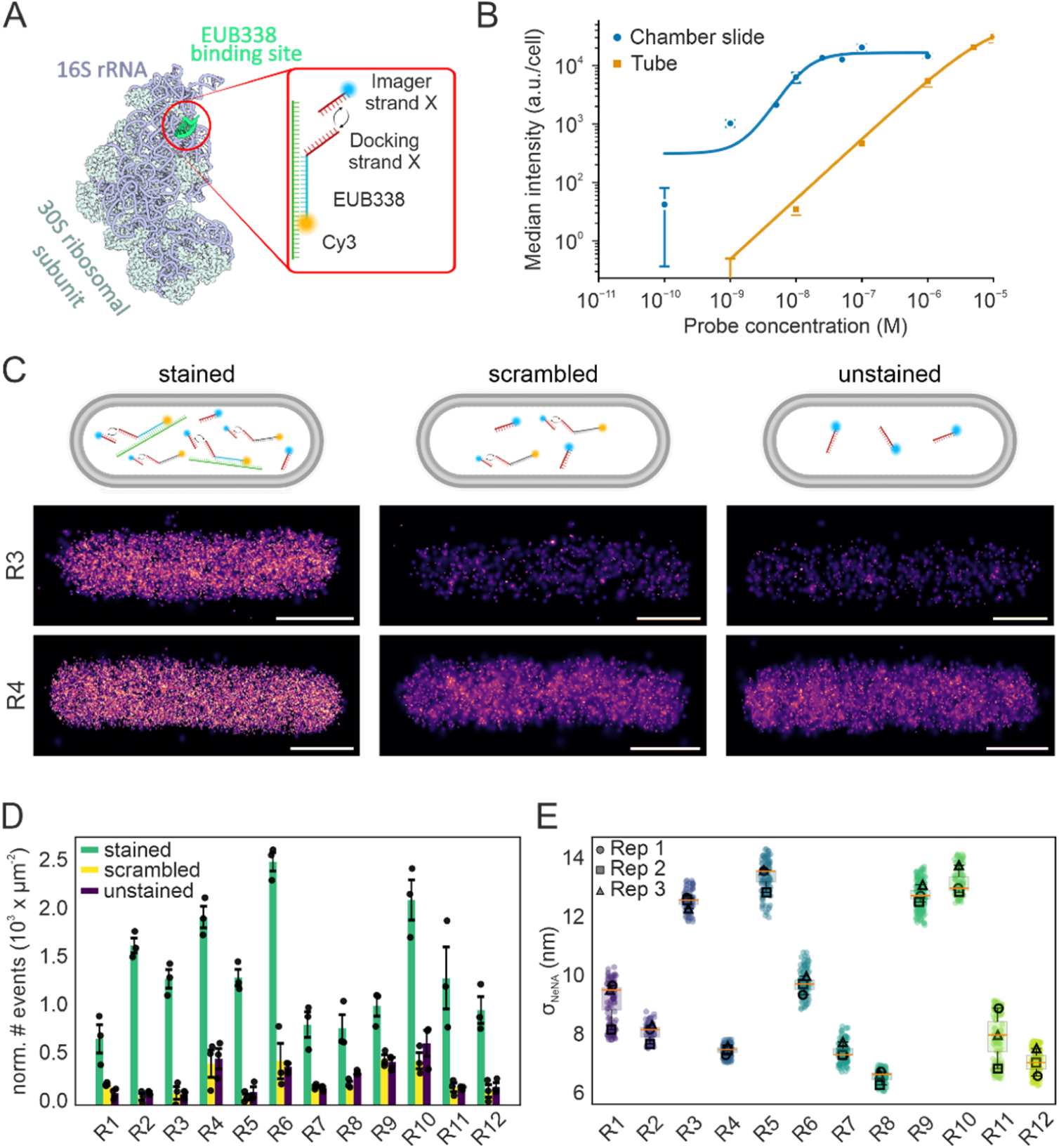
A platform for optimizing bacterial DNA-PAINT imaging. (A) Schematic of the rRNA-FISH strategy. The EUB338-probe binds specifically to an exposed loop of the E. coli 16S rRNA. The EUB338 probe carries a DNA-PAINT docking strand extension at the 3’ end, as well as a 5’ modification with Cy3. (PDB: 7OE1; rendered using 3dproteinimaging.com) [34,35]. (B) FISH workflow optimization. Staining of immobilized bacteria improves rRNA-FISH staining, as shown by probe titration using bacteria in solution or immobilized on chamber-slides. (C) Schematic of specificity control of EUB338-based screening and representative DNA-PAINT images of stained E. coli cells using EUB338-R3 and EUB338-R4. DNA-PAINT measurements of stained samples include specific signal, signal from unspecific binding of the FISH probe, and unspecific binding of the imager strand. The scrambled control gives insights on FISH-probe specificity, while the unstained sample reveals unspecific binding of the DNA-PAINT imager strand itself. (D) Comparison of normalized localization counts for imager strands R1-R12 under the conditions shown in (C). (E) Comparison of experimental localization precision using Nearest Neighbor Analysis (NeNA) [36]. D and E show the mean and SEM of three independent biological replicates. Scale bars in C are 1 µm. Scale bars are 1µm.

Importantly, the use of suitable controls reports on imager strand specificity, a parameter we found particularly important for high-quality DNA-PAINT imaging in bacteria. Using this system, we screened 12 imager-docking strand pairs [14,15] and optimized a subset of these sequences. We then used these sequences to demonstrate 4-plex Exchange-PAINT of intra- and extracellular targets in individual *E. coli* cells, as well as 4-plex Exchange-PAINT of effector mRNAs in *Y. enterocolitica*, thus providing a validated toolkit for super-resolution studies in Gram-negative bacteria. In summary, we present a simple, cost-efficient platform to evaluate the performance of known and future DNA-PAINT sequences in biological environments. The conserved nature of the EUB338 probe allows for easy transfer to other bacterial species and hopefully catalyzes broad adoption of DNA-PAINT imaging in bacteriology.

## Results

### An rRNA-FISH platform for optimizing DNA-PAINT imaging in a biological environment

We previously used RNA fluorescence *in situ* hybridization (RNA-FISH) in combination with DNA-PAINT to study mRNA distribution in the Gram-negative pathogen *Y. enterocolitica* using the first-generation DNA-PAINT sequence P1 [25]. When testing the next-generation, repetitive R-sequences[14,15], we experienced strong unspecific binding for some, but not all DNA-PAINT sequences. We hypothesized that the EUB338 probe, which we used as a positive control, could be a suitable system for testing the intrabacterial performance of different imager-docking strand pairs. The EUB338 probe has long been used for bacterial cell counting, as it offers a simple, low-cost readout of total ribosomal content [26,27]. Due to the high copy number of 16S rRNA (∼ 8,000 - 73,000 molecules per *E. coli* cell, depending on the generation time [28]), a single FISH probe is sufficient to generate a strong fluorescence signal. Next to determining bacterial load, EUB338 was employed to study ribosome heterogeneity [29,30] and for antibiotic profiling [31], demonstrating its versatility. By extending EUB338 with a DNA-PAINT docking sequence at the 3’ end, we adapted it as a reporter for ribosome nanoscale distribution. As FISH staining is highly sensitive to hybridization conditions [32], we added a Cy3 fluorophore to the 5’ end (Figure 1A). This provides, on one hand, a diffraction-limited readout on the staining intensity and serves as a staining control, while it, on the other hand, allows normalizing the DNA-PAINT signal across different EUB338-PAINT-sequence constructs. This dual-readout design allowed us to build a platform to benchmark candidate imager strands for bacterial DNA-PAINT imaging.

In our initial tests of the approach, we used the standard RNA-FISH protocol published by Skinner and colleagues, in which hybridization is performed in cell suspension [33]. We used a Cy5-EUB338 probe without docking site for direct readout and observed a strongly heterogeneous signal. Titrating the EUB338 probe resulted in better homogeneity, but saturation was not even reached at 10 µM probe concentration (Figure 1B). As homogeneous staining is essential for a reliable screen, we tested different staining protocols. We speculated that the high abundance of 16S rRNA depletes the FISH probe in suspension before saturation is reached. To reduce the biomass-to-probe ratio, we tested on-slide hybridization after immobilizing the fixed *E. coli* cells on Poly-L-lysine coated surfaces (Figure 1B). As this reduces the biomass-to-probe ratio by orders of magnitude, saturation was reached already at ∼200 nM concentration and the staining also resulted in a more homogeneous staining (Figure S1). We therefore used on-slide hybridization for all subsequent experiments.

### The EUB338 screening assay enables intra-bacterial benchmarking of imager–docking strand pairs

We have previously shown that extending EUB338 with a docking handle can be used for DNA-PAINT [25]. Having optimized the staining protocol, we started benchmarking different docking-imager strand pairs. We ordered FISH oligos that we designed with the 3’ sequence extension and a 5’-Cy3 modification. We selected the established R1 – R6 sequences [14], as well as the recently published R7 – R12 sequences [15] for our screen. We used 3 overlapping binding motifs to keep the overhang short and did not use spacers between the EUB338 and DNA-PAINT sequences. Readout of the Cy3 channel before DNA-PAINT imaging showed that different docking-strand extensions do not affect EUB338 hybridization efficiency (Figure S2). To test for non-specific signal and isolate its source, we included two negative controls: an unstained sample (no FISH probe added), reporting on non-specific binding of the imager strand itself and staining with a scrambled FISH probe, in which the EUB338 sequence was shuffled, while retaining the docking strand sequence. Signal detected in the latter control includes non-specific or off-target binding of the FISH probe, as well as unspecific binding of the imager strand. Comparison with the stained samples reveals the contribution of each source to the total signal (Figure 1C). We then acquired 3D DNA-PAINT data in biological triplicate using a biplane detection scheme [37] with epifluorescence illumination using imager strands at 25 – 100 pM concentration and a constant exposure time of 50 ms. The lack of optical sectioning required usage of ATTO655-conjugated imager strands, which enables DNA-PAINT imaging in epifluorescence illumination at concentrations in the picomolar to low nanomolar range. To facilitate efficient and reproducible analysis of the acquired DNA-PAINT data, we used a batch-analysis pipeline based on for segmentation, drift-correction, linking of localizations spanning multiple frames, and extraction of relevant parameters from the measurements (Fig S3, see Methods).

Unspecific binding varied substantially between imager strands (Figures 1C/D, S4), resulting in specific fractions of 60 - 92% (Specific fractions are defined as 1-(events_unstained_/events_stained_), referred to as specificity from here on). R2 and R3 showed the highest specificity (92.0 ± 0.1% (SEM) and 91.6 ± 1.5%, respectively), while R4 and R8 showed substantially more non-specific signal (74 ± 6% and 61 ± 7%, respectively). Of note, a very conservative threshold was used for single-molecule localization to account for differences in brightness between the different docking-imager strands pairs. Scrambled and unstained controls resulted in similar localization counts per area (determined via segmentation) for most imager strands, indicating that non-specific binding is driven predominantly by the imager strand itself rather than by non-specific binding or off-target FISH-probe hybridization. Due to the high abundance of 16S RNA in fast-growing *E. coli* cells, the imager strand concentrations used here were rather low, and imaging of low-abundance targets can require concentrations up to a few nM. We thus tested the imager strands also at a higher concentration of 1nM, resulting in overlapping binding events too dense to analyze. Nevertheless, it revealed a substantial amount of unspecific binding for the R2 imager strand and even more for R4 (Figure S5). R7–R12 strands showed systematically lower specificity (e.g. R12: 77.9%) under the initial screening conditions (Table 1).

**Table 1:** Performance characterization of different imager strands in E. coli cells. Different imager strand lengths are indicated after underscores. Identical acquisition and analysis parameters were chosen for better comparability (50 ms, SBR 15 for specificity, σ_NeNA_, photon count and localization precision; 20 ms and 10 pM concentration for determination of kinetic parameters). #Photons represents the integrated photon count after linking of subsequent-frame localizations. Errors represent SEM and were determined from mean values of biological triplicates.

| Imager strand | Specificity ± SEM | $\sigma_{\text{NeNA}} \pm$ SEM (nm) | #photons ± SEM | $\sigma_{\text{XY}} \pm$ SEM (nm) | $\sigma_z \pm$ SEM (nm) | binding time ± SEM (ms) | $K_{\text{off}} \pm$ SEM ($\text{s}^{-1}$) |
| --- | --- | --- | --- | --- | --- | --- | --- |
| R1 | $0.80 \pm 0.08$ | $9.5 \pm 0.6$ | $2120 \pm 300$ | $11.1 \pm 0.7$ | $37.9 \pm 1.2$ | $180 \pm 14$ | $6.0 \pm 0.3$ |
| R2_6nt | $0.920 \pm 0.001$ | $7.9 \pm 0.3$ | $4010 \pm 70$ | $7.2 \pm 0.2$ | $25.7 \pm 3.3$ | $264 \pm 15$ | $4.0 \pm 0.2$ |
| R2_7nt | $0.869 \pm 0.004$ | $8.0 \pm 0.2$ | $5100 \pm 40$ | $6.1 \pm 0.1$ | $18.0 \pm 0.2$ | $418 \pm 17$ | $2.6 \pm 0.1$ |
| R3 | $0.92 \pm 0.02$ | $12.7 \pm 0.2$ | $1700 \pm 50$ | $14.0 \pm 0.2$ | $41.0 \pm 0.6$ | $287 \pm 16$ | $3.4 \pm 0.4$ |
| R4_6nt | $0.7 \pm 0.1$ | $7.7 \pm 0.2$ | $5200 \pm 60$ | $6.8 \pm 0.3$ | $26.6 \pm 2.1$ | $306 \pm 15$ | $2.9 \pm 0.2$ |
| R4_7nt | $0.97 \pm 0.01$ | $8.2 \pm 0.1$ | $4400 \pm 60$ | $6.9 \pm 0.1$ | $20.9 \pm 0.3$ | $422 \pm 26$ | $2.1 \pm 0.1$ |
| R5_7nt | $0.89 \pm 0.04$ | $13.5 \pm 0.2$ | $1600 \pm 50$ | $15.9 \pm 0.5$ | $54.2 \pm 6.2$ | $210 \pm 27$ | $5.6 \pm 0.6$ |
| R5_8nt | $0.989 \pm 0.001$ | $12.0 \pm 0.5$ | $2200 \pm 20$ | $12.4 \pm 0.1$ | $39.6 \pm 0.7$ | $322 \pm 29$ | $2.8 \pm 0.4$ |
| R5_9nt | $0.946 \pm 0.003$ | $10.4 \pm 0.4$ | $2400 \pm 50$ | $11.5 \pm 0.2$ | $36.5 \pm 0.3$ | $600 \pm 80$ | $1.9 \pm 0.3$ |
| R6 | $0.84 \pm 0.03$ | $9.9 \pm 0.4$ | $4000 \pm 400$ | $7.8 \pm 0.4$ | $22.4 \pm 1.1$ | $188 \pm 16$ | $5.8 \pm 0.3$ |
| R7 | $0.77 \pm 0.06$ | $8.1 \pm 0.6$ | $3030 \pm 210$ | $9.5 \pm 0.6$ | $33.1 \pm 2.7$ | $144 \pm 8$ | $7.1 \pm 0.3$ |
| R8 | $0.6 \pm 0.1$ | $6.9 \pm 0.3$ | $4400 \pm 150$ | $6.4 \pm 0.2$ | $20.4 \pm 0.4$ | $431 \pm 23$ | $2.4 \pm 0.2$ |
| R9 | $0.6 \pm 0.1$ | $13.3 \pm 0.2$ | $1300 \pm 60$ | $19.9 \pm 0.2$ | $59.7 \pm 3.1$ | $230 \pm 25$ | $4.9 \pm 0.6$ |
| R10 | $0.7 \pm 0.1$ | $13.3 \pm 0.4$ | $1900 \pm 90$ | $14.2 \pm 0.2$ | $42.3 \pm 2.3$ | $138 \pm 5$ | $7.3 \pm 0.3$ |
| R11 | $0.8 \pm 0.1$ | $8.2 \pm 0.7$ | $4040 \pm 80$ | $7.8 \pm 0.3$ | $25.7 \pm 1.7$ | $330 \pm 120$ | $4.1 \pm 1.7$ |
| R12 | $0.8 \pm 0.1$ | $7.2 \pm 0.3$ | $4400 \pm 250$ | $6.6 \pm 0.4$ | $19.6 \pm 0.5$ | $462 \pm 25$ | $1.9 \pm 0.2$ |

To estimate the spatial resolution of each docking-imager strand pair, we determined experimental localization precision via Nearest-Neighbor Analysis (NeNA) [36]. In contrast to the theoretical localization precision, NeNA provides an experimental estimate of localization precision from distances between localizations in consecutive frames (from here on referred to as σ_NeNA_). σ_NeNA_ values for the different docking-imager strand pairs ranged between 6.8 ± 0.3 (R8) and 13.5 ± 0.2 nm (R5) (Figure 1E). Imager strands composed only of G and T bases (R2, R4, R6) in general achieved better (lower) σ_NeNA_ values than GA-based imagers (R1, R3, R5), consistent with their higher photon budget (Figure S6). However, this might be a dye-specific effect, which might be circumvented by acquisition in TIRF or HILO mode using other fluorophores [38]. However, as we used a comparably fast exposure time, splitting of the signal into several imaging frames results in larger σ_NeNA_ values, reflecting poorer per-frame localization precision than would be obtained by integrating photons over the complete binding event.

We next wanted to determine the hybridization kinetics of DNA-PAINT docking-imager strand pairs within tightly packed bacterial cells. DNA-PAINT acquisition consists of alternating bright phases (τ_ON_ or ON-time; imager strand bound to the docking strand) and dark phases (τ_OFF_ or dark time; free docking strand), characterized by the dissociation rate k_off_ (≈ 1/τ_ON_) and apparent association rate k_on_ (≈ 1/(τ_OFF_ × [imager])), respectively. Both rates are sensitive to imaging buffer composition (primarily salt concentration), temperature, and imager–docking sequence design [19,39].

To resolve individual binding events and reliably calculate τ_ON_, we reduced the imager strand concentration to 10 pM and shortened the exposure time to 20 ms. This resulted in τ_ON_ distributions that were best approximated by a bi-exponential decay function. Fitting revealed two populations, a long-dwell population consistent with specific imager strand binding to the docking strand, and a short-dwell population consistent with transient, non-specific interactions (Figures 2A and S7A). This assignment was validated by measuring unstained controls, which should only report on non-specific interactions. Indeed, the ON-times of the unstained controls agree well with the short-dwell component determined in our DNA-PAINT experiments across imager strands (Figure S7B). k_off_ per cell was calculated from the inverse of the long-dwell ON-time (Figure S7C). All kinetic parameters for the different imager strands are summarized in Table 1.

**Figure 2:**
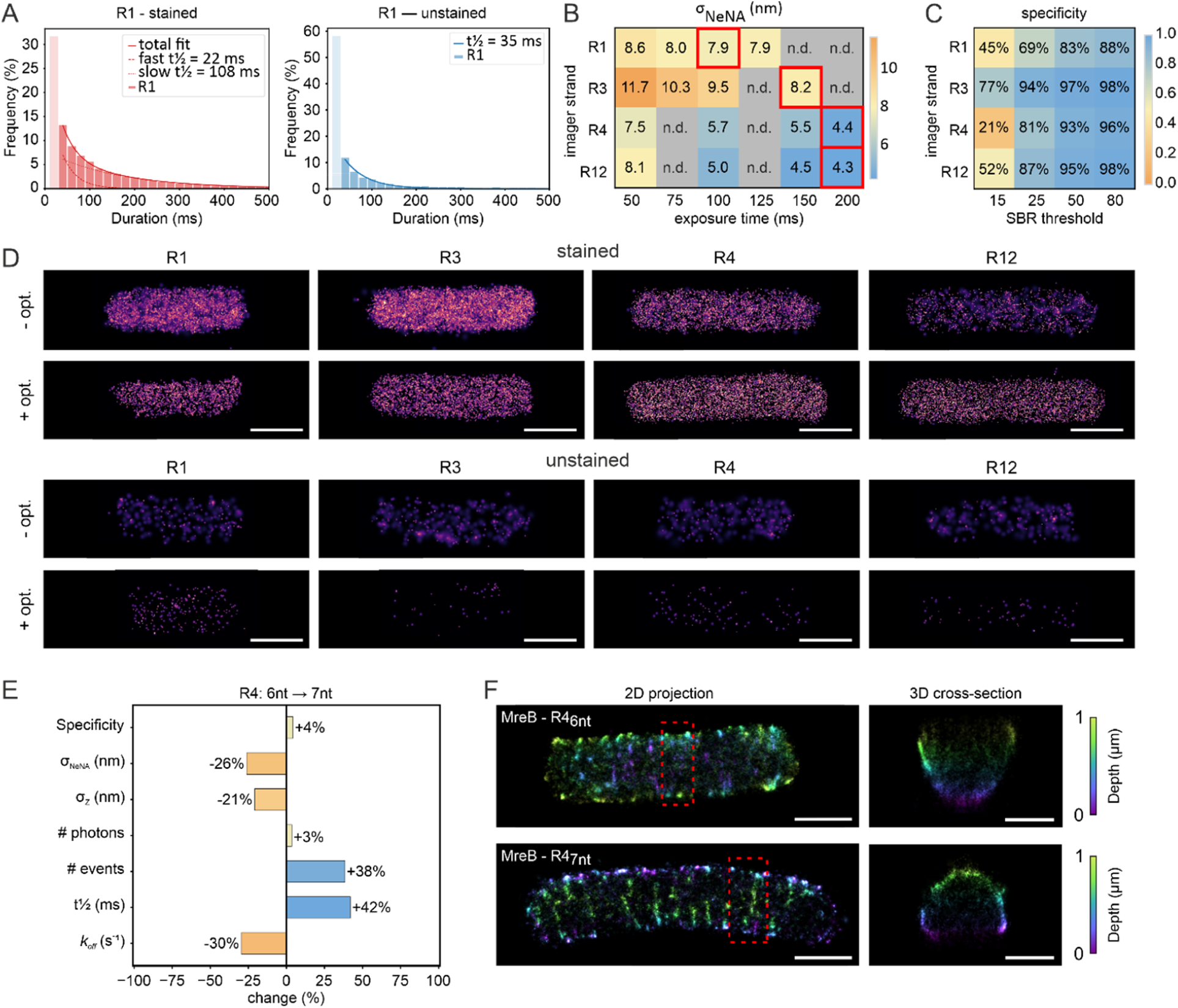
Optimization of DNA-PAINT imaging based on kinetic binding analysis. (A) ON-time analysis for R1 docking-imager strand pair in stained samples and unstained controls. Fitting with a bi-exponential decay function reveals the presence of a long-dwell and short-dwell fraction in stained samples, while only the short-dwell fraction is present in the unstained control. (B) Heatmap comparing tested exposure times and the extracted σ_NeNA_ values for imager strands R1, R3, R4_7nt_ and R12. (R1 = 84 cells; R3 = 26 cells; R4 = 25 cells; R12 = 29 cells) (C) Heatmap comparing the specificity to the SBR threshold used for single-molecule localization recorded using optimized exposure times. The selected exposure times are indicated by red squares in (B). (D) Representative cells recorded under suboptimal and optimized DNA-PAINT conditions. Optimal conditions were 100 ms for R1, 150 for R3, 200 ms for R4_7nt_ and 200 ms for R12 using a SBR of 80 for all sequences. (E) Relative changes for extension of R4_6nt_ to R4_7nt_ variants using optimized imaging and analysis parameters. (R4_6nt_ = 29 cells; R4_7nt_ = 23 cells) (F) Representative images of MreB^sw^-sfGFP acquisitions using R4_6nt_ and R4_7nt_ imager strands under optimized conditions. 30,000 frames were recorded with 200 ms exposure and at 250 pM imager-concentration. Scale bar grown under secretion-inducing conditions are 1 µm for the 2D projection in (D) and 600 nm for 3D cross-sections in (F).

### The EUB338 platform enables *in situ* optimization of DNA-PAINT imaging parameters

Our screening of DNA-PAINT sequences provided insights into imager strand specificity and kinetics. However, as we used a fixed, fast exposure time and analysis parameters, we likely imaged each docking-imager strand pair under suboptimal conditions. Indeed, our ON-time analysis revealed binding half times in the range of 140 – 460 ms, and even longer times for extended binding sequences (e.g. 600 ms for R5 imager strand with 9 nucleotides). These times differ substantially from the 50 ms exposure time used in the initial screen. As NeNA values are calculated on distances of subsequent-frame localizations, distribution of the single-molecule signal on many frames leads to decreased per-frame brightness and thus higher σ_NeNA_ values compared to acquisitions with optimized exposure times. Adjusting exposure times to the average ON-time should thus lead to improved localization precisions. We tested the effect of exposure time on four imager strands spanning a range of specificity, kinetics, and localization precisions (R1, R3, R4, R12). The exposure times were selected based on the ON-time measured for each imager strand (Figures 2A and S7). We then determined σ_NeNA_ values for each exposure time, which revealed different optimal values for the imager strands analyzed (Figure 2B). We found an exposure time of 100 ms to be ideal for R1, 150 ms for R3, and 200 ms for R4 and R12, broadly reflecting the differences in determined ON-times (Table 1). This is well expected, as increasingly matching exposure times allow the collection of more photons until they reach or exceed the ON time of the imager strand. However, longer exposure times also integrate more background signal, which can increase the number of false-positive localizations if detection thresholds are not adjusted. To determine the localization threshold providing the best specificity, we re-analyzed the same measurements with a range of signal-to-background (SBR) thresholds during the localization step (Figure 2C). Specificity increased with increasing thresholds for all imagers, reaching 88% for R1 and 96-98% for R3, R4 and R12 using a SBR threshold of 80.

Because the 7 nt variant of R4 showed a longer ON-time than the 6 nt variant (Table 1), we compared them directly in EUB338-R4-stained cells using the optimized acquisition settings established above (200 ms exposure, SBR threshold 80). Extending the imager strand length to 7 nt improved both specificity and σ_NeNA_ in comparison to the 6 nt variant (Specificity of 88.39% and 95.67%, σ_NeNA_ of 5.7 ± 0.5 nm and 4.3 ± 0.3 nm for R4_6nt_ and R4_7nt_, respectively), although at the expense of longer acquisition times (Figure 2E).

To test whether this improvement also allows for improved imaging of a biological target, we compared the R4_6nt_ and R4_7nt_ imager strand variants on MreB, a bacterial actin homolog that forms filamentous structures at the cytosolic side of the inner membrane. MreB is a great bacterial model structure for super-resolution imaging, as it forms antiparallel double filaments with a defined width of approximately 6–7 nm based on the structurally characterized *Caulobacter crescentus* MreB double filament. [40]. We used a strain expressing an MreB^sw^-sfGFP fusion [41] that we labeled with a cocktail of two anti-GFP nanobodies, both carrying an R4-docking strand (Figure 2F). Both R4_6nt_ and R4_7nt_ revealed MreB filaments, however, R4_7nt_ yielded fewer localizations in the cytosol compared to R4_6nt_, confirming the increase in specificity observed for EUB338. In agreement with EUB338 experiments, σ_NeNA_ also strongly improved from 15.1 ± 7.7 nm for R4_6nt_ to 7.5 ± 1.3 nm for R4_7nt_, improving the experimental lateral localization precision approximately twofold, while also affecting other parameters (Figure 2E). Both effects are evident in the 3D reconstructions, which provide a higher image quality for R4_7nt_ under optimized imaging and analysis conditions (Figure 2F, Supplementary Video 1).

### 3D Exchange-PAINT provides insights into the cellular architecture of *E. coli* cells

Having optimized the imaging and analysis parameters for selected DNA-PAINT sequences, we aimed to visualize bacterial targets in and on individual *E. coli* cells using 3D Exchange-PAINT. To probe different cellular compartments, we selected surface-exposed lipopolysaccharides (LPS, outer leaflet of the outer membrane), MreB (anchored to the cytosolic side of the inner membrane), RpoC, the β′ subunit of RNA polymerase (cytosolic) and the DNA, the latter recorded with a conventional PAINT approach using JF_646_-Hoechst [42]. Imaging under optimized conditions provided high-quality 3D (DNA-)PAINT images, for which we achieved mean σ_NeNA_ values of 6.3 ± 0.2 nm (DNA), 5.7 ± 0.6 nm (LPS), 6.5 ± 0.4 nm (MreB) and 7.0 ± 1.0 nm (RpoC), and mean axial localization precisions of 16.9 ± 0.4 nm (DNA), 24.2 ± 1.7 nm (LPS), 27.1 ± 1.2 nm (MreB) and 23.9 ± 0.9 nm (RpoC) (Figure 3A, Supplementary Video 2).

**Figure 3:**
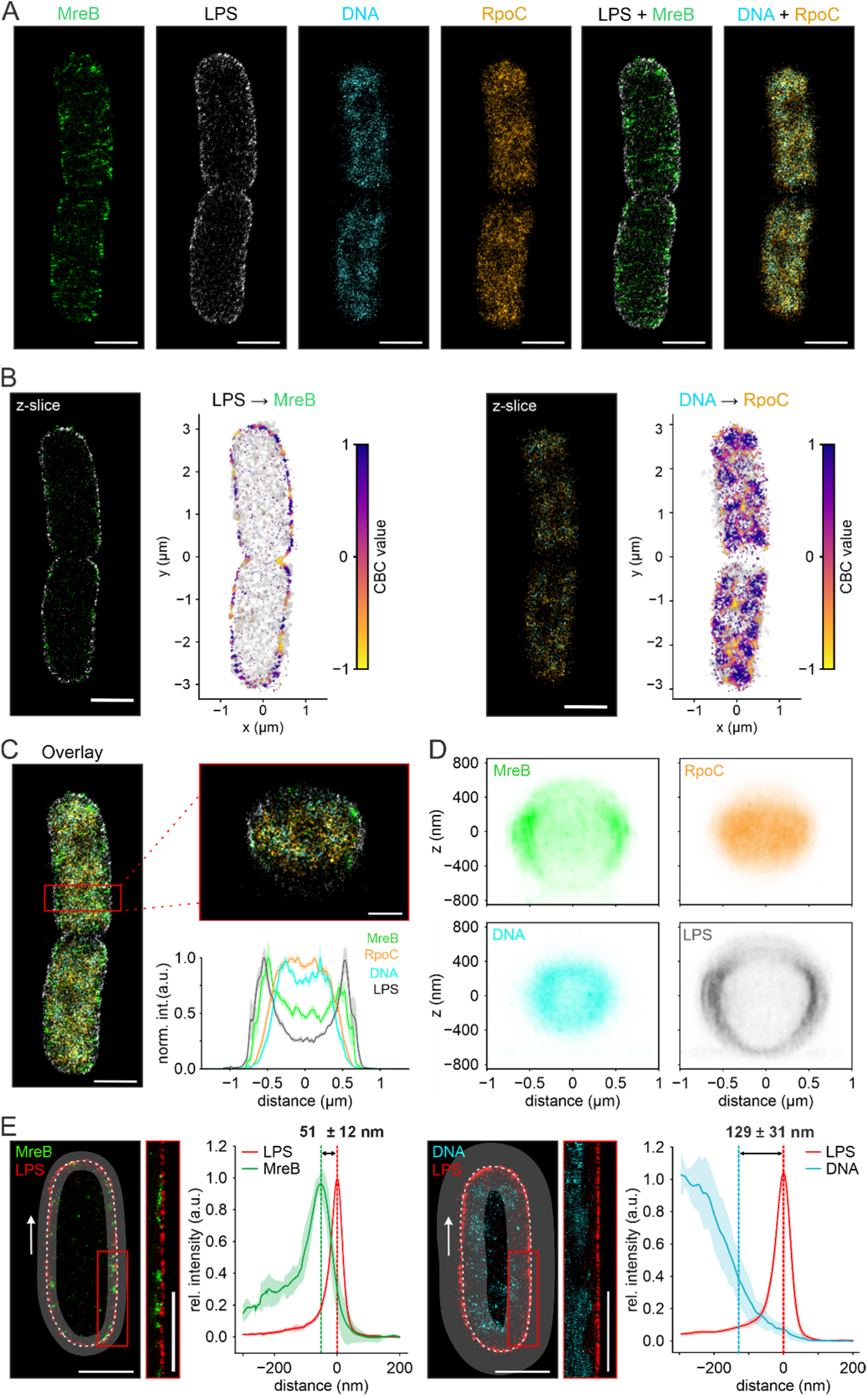
Exchange-PAINT of protein targets in E. coli using optimized imaging conditions. (A) Multi-target DNA-PAINT/PAINT images of E. coli cells fixed in exponential phase under fast growing conditions. Shown is the 100 nm center z-section of MreB^sw^-sfGFP (labeled with @GFP nanobodies conjugated to R1, green) LPS (anti-E. coli antibody labeled with a secondary probe conjugated with R12, grey), RpoC (RpoC-ALFA, anti-ALFA nanobody [50] conjugated with R3, orange) and DNA (JF_646_-Hoechst, cyan), as well as selected overlays. For the representative cell shown, single-cell σ_NeNA_ values of 5.1 nm (LPS), 6.7 nm (MreB), 8.3 nm (RpoC) and 6.6 nm (DNA), and an axial localization precision of 24.9 nm, 27.2 nm, 23.7 nm and 16.7 nm were determined. (B) Coordinate-based colocalization (CBC) analysis [47] using a 200 nm z-slice of the cell shown in (A). CBC score rendering of LPS localizations relative to MreB (left) and RpoC localizations relative to DNA (right). Scores range from −1 (anticorrelated localization) to +1 (fully colocalized). (C) Four-color overlay of MreB, RpoC, DNA and LPS. The red box marks the region of the cross-sectional view. The transverse line profile (mean ± SD) shows the probability density of each target across the cell width. (D) Average cross-sectional distribution of the 4 targets (N_cells_ = 14). Individual cells were aligned using custom code (see methods). (E) Distance distribution of MreB and LPS using a straightening approach. Signal along the cell perimeter was extracted from the straightened membrane contour (top, scale bar: 1 µm). The mean relative intensity profile (± SD) shows that the MreB and LPS maxima are separated by 51 ± 12 nm (N_cells_ =13). The same analysis for membrane and DNA provided a distance of 129 ± 31 nm between the LPS distribution maximum and 1/e cutoff for the DNA distribution (N_cells_=13). Scale bars are 1µm in (A), (B), (C, 2D projection) and (E), and 400 nm in (C) for the 3D cross-section.

Each target showed its expected subcellular localization (Figure 3A). LPS formed a continuous layer coating the outer membrane, with few localizations in the cell cytosol. For MreB, we observed filaments and small patches as reported in literature, being mostly located at the inner membrane along with a smaller population of cytosolic localizations. Whether the cytosolic localizations reflect non-specific imager binding, non-specifically bound nanobodies or a genuine cytosolic pool of monomeric MreB [43] needs to be determined. RpoC showed a nucleoid-like distribution that is expected under the growth condition used, where ∼85% of RNA polymerase complexes were reported to be engaged in active transcription [44]. This is visually verified in the overlay of RpoC and DNA channels, which shows strong signal overlap. The DNA itself exhibited the bilobed nucleoid organization previously reported for fast-growing cells [45,46].

We then applied coordinate-based colocalization (CBC) [47] (see Methods) to test the spatial correlation between different targets. MreB and LPS localizations showed positive colocalization scores (CBC = 0.3 ± 0.2; Figures 3B and S8), consistent with the close proximity of outer and inner membrane in *E. coli*. Similarly, RpoC and DNA showed a high colocalization fraction concentrated over the DNA lobes (CBC = 0.62 ± 0.08; Figures 3B and S8), consistent with the substantial DNA-bound RpoC population described above. A transverse line intensity profile across the cell confirmed clear spatial separation between the cytosolic markers (RpoC, DNA) and the membrane markers (MreB, LPS) (Figure 3C).

As our biplane data provides 3D information, we can also access target distributions in cylindrical cross-sections. Using a custom-written code (see Methods), we aligned the cross-sections of the cylindrical parts of 14 cells and generated a map reporting on the targets’ average axial distributions (Figure 3D). This analysis revealed the expected localizations for LPS and MreB at the cell membranes, as well as the membrane-directed, and center-devoid localization of the nucleoid reported recently [48,49] (Figure 3D, Supplementary Video 2).

Despite the colocalization observed in our CBC analysis, we noted a small, consistent gap between the LPS and MreB signals, reflecting separation of outer and inner membranes by the periplasmic space. We thus wondered whether the high resolution of DNA-PAINT allows to quantify distances between the Gram-negative cell membranes. Having LPS and MreB as markers for the inner and outer membrane, we extracted the central slice of each cell, straightened it along the LPS-defined perimeter, measured the signal distributions by line plotting and extracted peak-to-peak distances between LPS and MreB signals from the resulting intensity profiles (Figures 3E and S9). Our analysis reported a mean distance of 51 ± 12 nm (N_cells_ = 13 cells). This value is in line with expectations based on literature values of the periplasm width (13 - 25nm, [51,52]) when accounting for the linkage errors introduced by the labeling strategy (∼15-20 nm, combined for both labels), the localization precision, as well as the label positioning on opposing sites of ∼ 5 nm wide membrane lipid bilayers. Based on this, we estimate a periplasmic width of ∼20-30 nm for *E. coli* cells during exponential growth in rich medium.

Together, these analyses of our 4-color Exchange-PAINT experiment demonstrate that the high resolution, obtained under optimized conditions, can unlock relevant biological information regarding cellular architecture and the distribution of different biomolecules.

### 3D Exchange-PAINT unravels the spatial correlation of pathogen effector mRNAs

Understanding the molecular mechanisms of bacterial infection remains critical in the context of rising antibiotic resistance. One key virulence mechanism is the type III secretion system (T3SS), a multi-protein injection apparatus that translocates effector proteins from the bacterial cytosol directly into host cells. These effectors have various functions, such as suppression of the host immune system and facilitation of pathogen uptake [53,54]. The Gram-negative pathogen *Yersinia enterocolitica* uses a T3SS encoded on its virulence plasmid (pYV) to deliver Yop (*Yersinia* outer protein) effectors into host cells [55]. We recently investigated whether the YopM effector protein is secreted via a spatiotemporal coupling of transcription, translation and insertion (short: transertion) into the T3SS. In this case, one would expect the mRNA encoding the effector in close proximity to the bacterial inner membrane. As *yopM* transcripts showed a purely cytosolic distribution, transertion is likely not involved in YopM secretion [25]. Given the high multiplexing capability of DNA-PAINT, we wanted to extend this study to other effector proteins, and we visualized the three effector mRNAs *yopM*, *yopE*, and *yopH* under different conditions. This included a condition which induces secretion in wild type (WT) *Y. enterocolitica* cells *in vitro* (secretion-inducing), growth of the WT under non-secreting conditions, as well as a strain lacking the plasmid of *Yersinia* virulence (pYV ^−^) not expressing any T3SS components. The pYV ^−^ strain serves as negative control under secretion-inducing conditions and can be compared to the scrambled control of our EUB338 platform. Hence, it reports on the non-specific background and allows quantification of WT mRNA expression under secreting- and non-secreting conditions.

3D Exchange-PAINT resolved distinct mRNA clusters in secreting WT cells for all three transcripts, while both negative controls showed markedly fewer localizations (Figure 4A, Supplementary Video 3). Signal in the pYV ^−^ control reflects non-specific probe/imager background; the residual signal in uninduced WT is consistent with basal transcription in the absence of induction [57]. To illustrate the method’s ability to quantify spatial relationships between targets, we performed CBC analysis between all three mRNA pairs (Figure 4B). The analysis showed a slight colocalization (Mean CBC = 0.15) (Figure S10) for the effector mRNA signals, indicating that they localize to the same subcellular area (cytosolic with nucleoid exclusion) [25]. To get an estimate of transcript levels, we determined the number of localization clusters per cell via 3D-DBSCAN analysis (see Methods). We found that *yopH* and *yopE* showed an average number of 77 ± 24 and 67 ± 21 clusters per cell under secreting conditions, clearly above both negative controls (pYV^−^: *yopE* 7 ± 4, *yopH* 13 ± 5 cluster per cell; non-secreting WT: *yopE* 4 ± 3, *yopH* 17 ± 7 cluster per cell) (Figure 4C). *yopM* showed fewer clusters overall (40 ± 17 cluster per cell), alongside with a higher background level in its negative controls (non-secreting 16 ± 9, ΔpYV 19 ± 10 cluster per cell). This finding is consistent with lower *yopM* transcript abundance reported for the secretion time point and agrees well with the numbers that we reported previously [25]. Alternative explanations are less specific FISH probes or the different performance of the imager strand sequence.

**Figure 4:**
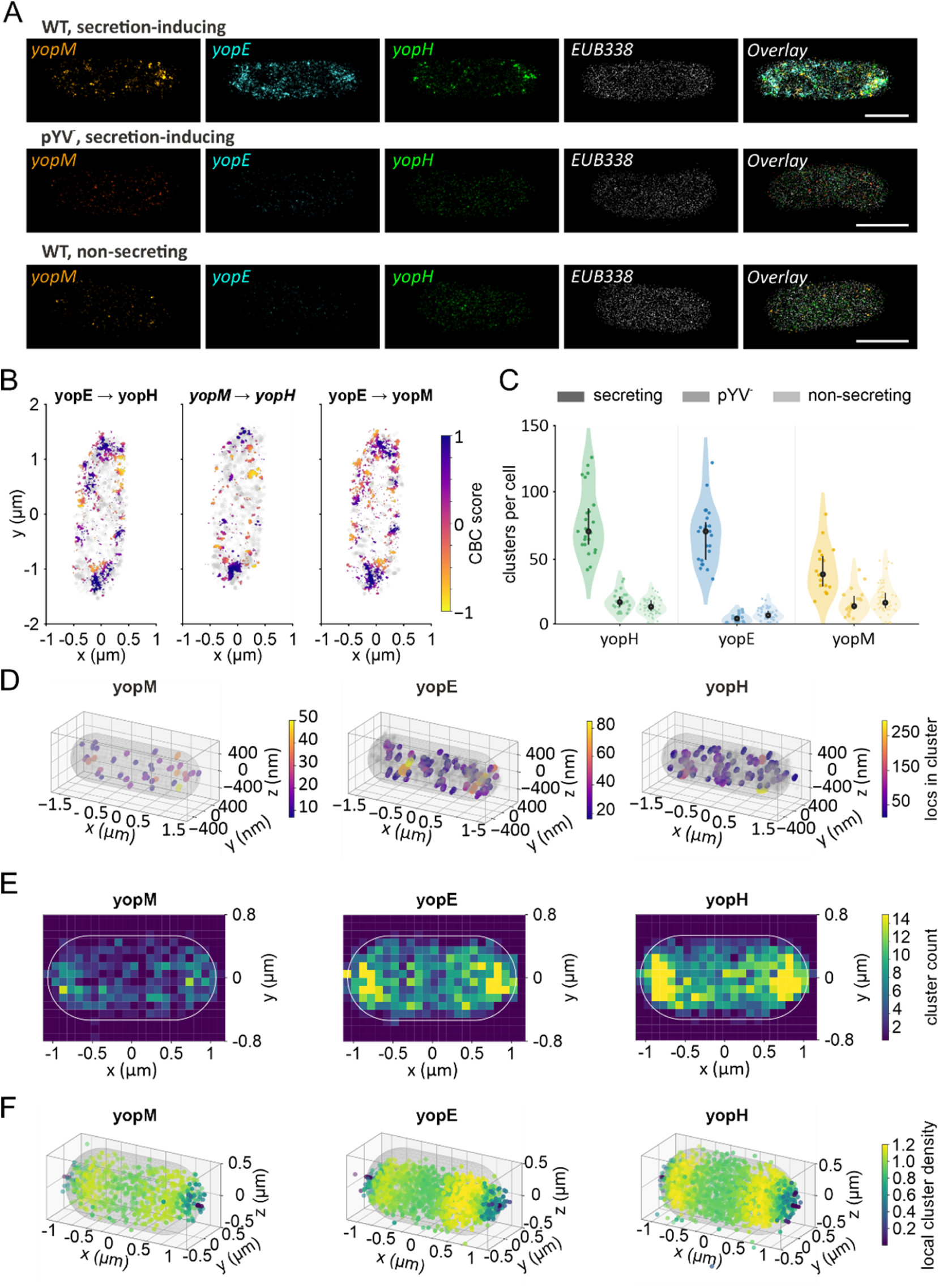
Spatial organization of T3SS effector mRNAs in Yersinia enterocolitica under different conditions. (A) Multiplexed DNA-PAINT images of yopM (orange), yopH (green) and yopE (cyan) mRNA labeled via RNA-FISH, with EUB338 (grey) as reference for the cell outlines. Cells were grown under secretion-inducing (WT and pYV^−^) and non-secreting (WT) conditions. Overlays of all channels are shown on the right. Scale bars: 1 µm. (B) CBC analysis of the induced WT cell shown in (A): yopM relative to yopE, yopM relative to yopH, and yopE relative to yopH. Scores range from −1 (anticorrelated signal) to +1 (fully colocalized signal). (C) Quantification of yopH, yopE and yopM localization clusters per cell using 3D-DBSCAN under non-secreting (WT) and secreting (WT and pYV^−^ strains) conditions. Data points represent individual cells; black markers show mean ± SD. (D) 3D representation of detected clusters of representative cells, colored by the number of localizations per cluster (left to right: yopM, yopE, yopH) (code adapted from [56]). Gray area represents a 3D cell model calculated on EUB338 signal. (E) Pooled cluster density maps of secreting cells for yopM, yopE and yopH, 2D projected onto the long and short cell axes. The white outline indicates the average cell shape (N_cells_ = 14, single replicate). (F) 3D local cluster density of pooled clusters for yopM, yopE and yopH in secreting cells. (N_cells_=14, single replicate).

To further resolve the spatial distribution of RNA-FISH clusters within the sphero-cylindrical cell space, we computed the 3D distance of each cluster to the cells’ center of mass. We generated both 3D cluster maps (Figure 4D) and heatmaps of cluster density (Figure 4E) along the cell’s long and short axes and in 3D space (Figure 4F). All three effector mRNAs showed clear accumulation in the cytoplasmic space directed towards the cell poles (see also Supplementary Video 3), illustrating the method’s ability to resolve sub-diffraction-limit spatial patterns of individual transcript populations. Clusters in the negative controls, by contrast, were more distributed towards the cell center (Figure S11). Because our cell-alignment method does not distinguish old from new pole, the current approach cannot resolve polarized transcript accumulation.

Of note, this analysis is based on a single biological replicate (see Figure 4C caption for cell counts) and is intended to demonstrate the capability of Exchange-PAINT to resolve individual mRNA clusters at high spatial resolution and in 3D, rather than to establish new quantitative biology.

## Discussion

In this study, we present a generalizable *in situ* framework for benchmarking, optimizing, and applying DNA-PAINT docking-imager strand pairs to 3D super-resolution imaging of Gram-negative bacterial cells. Imager strands for bacterial DNA-PAINT imaging have so far been adopted from DNA-origami or mammalian-cell studies [14,19] without systematic benchmarking, even though bacteria provide a distinct environment: a small cytoplasmic volume, a cell envelope that constrains probe accessibility, and strong molecular crowding. The EUB338-based FISH-PAINT screen closes this gap by using an abundant, well-characterized target (16S rRNA) as an internal calibration standard [26]. The simple extension of the EUB338 probe with docking sequences, the need for only a single FISH probe and the high level of control using a scrambled sequence, allows to generate a robust and straightforward approach to quantitatively assess performance of DNA-PAINT sequences *in situ*.

Our screening of 12 repetitive DNA-PAINT sequences (R1 to R12) [14,15] on a biplane 3D system provided several insights. First, the use of consistent acquisition and analysis parameters revealed differences in the photon flux (number of photons per frame) of ATTO655-conjugated imager strands (Table 1), causing up to 2.5-fold differences in localization precision (R8: σ_NeNA_ = 6.9 ± 0.3 nm; σ_xy_ = 6.4 ± 0.2 nm; σ_z_ = 20.5 ± 0.5 nm; vs. R5: σ_NeNA_ = 13.5 ± 0.2 nm; σ_X_ = 15.9 ±0.5 nm; σ_Z_ = 54 ± 6 nm) (Figure 1E) and specificities ranging from 60 – 92% for non-extended sequences (Table 1). These differences can be partly explained by the choice of dye, as imaging in epifluorescence forced us to use ATTO655, which is known to be quenched by certain nucleobases [58]. Second, ON-time analysis using imaging with short exposure times and low imager strand concentrations revealed that the varying imager strand specificity can be explained by differences in unspecific binding. Here, the ON-time distribution could be fitted well with a bi-exponential decay function, with the fast species matching the half-time of the unstained controls (Figures 2A and S7). As expected, increasing imager strand length for selected sequences by 1 or 2 nucleobases resulted in longer binding times (Table 1). This can improve the resolution by collecting more photons, but results in longer acquisition times. Based on the ON-time analysis results, we performed a guided parameter sweep by adjusting exposures times and fitting parameters for DNA-PAINT sequences R1, R3, R4 and R12. These optimizations strongly improved specificity and σ_NeNA_ values (e.g. from 7.2 ± 0.3 nm to 4.0 ± 0.8 nm for R12), demonstrating the strength of our approach in optimizing DNA-PAINT imaging. It also revealed that extending imager strand length can transform a low-performing into a well-performing sequence. We observed that extending R4 imager strand length from 6 nt to 7 nt improved both the specificity from 88.4% to 95.7% and σ_NeNA_ values from 5.7 ± 0.5 nm to 4.3 ± 0.3 nm. The effect on 3D image quality becomes evident when testing both variants on protein targets. Using the filament-forming actin homologue MreB as a model target, the increase in precision (15.1 ± 7.7 nm for R4_6nt_ vs 7.5 ± 1.3 nm for R4_7nt_) and specificity enhanced the visualization of the membrane-associated filaments (Figure 2F, Supplementary Video 1), demonstrating that the results from EUB338 probe can be transferred to protein targets (Table 2).

**Table 2:** Summary of parameters of optimized imager strand conditions of EUB338 and cellular targets. N_cells_ = 14 (MreB-R1, RpoC-R3 and LPS-R12), 74 (EUB338-R1), 2S (EUB338-R3), 2C (EUB338-R12) and 2C (EUB338-R47nt). N_replicates_ = 1.

| Target / Imager strand | Specificity ± SD | $\sigma_{\text{NeNA}} \pm$ SD (nm) | # Photons ± SD | $\sigma_{\text{xy}} \pm$ SD (nm) | $\sigma_z \pm$ SD (nm) |
| --- | --- | --- | --- | --- | --- |
| EUB338-R1 | $0.88 \pm 0.02$ | $6.4 \pm 0.4$ | $6880 \pm 130$ | $5.9 \pm 0.2$ | $18.7 \pm 0.6$ |
| MreB-R1 | $0.85 \pm 0.03$ | $6.5 \pm 0.4$ | $6370 \pm 100$ | $7.5 \pm 0.3$ | $27 \pm 2$ |
| EUB338-R3 | $0.98 \pm 0.01$ | $7.7 \pm 0.4$ | $6090 \pm 110$ | $7.1 \pm 0.2$ | $22 \pm 2$ |
| RpoC-R3 | $0.93 \pm 0.03$ | $7.0 \pm 1.0$ | $8600 \pm 2500$ | $6.7 \pm 0.9$ | $24 \pm 2$ |
| EUB338-R12 | $0.98 \pm 0.01$ | $4.0 \pm 0.8$ | $20500 \pm 11800$ | $3.1 \pm 0.5$ | $12 \pm 1$ |
| LPS-R12 | $0.83 \pm 0.04$ | $5.7 \pm 0.6$ | $11400 \pm 400$ | $5.7 \pm 0.3$ | $24 \pm 2$ |
| EUB338-R4<sub>7nt</sub> | $0.96 \pm 0.02$ | $4.3 \pm 0.3$ | $15800 \pm 2000$ | $3.5 \pm 0.7$ | $13 \pm 10$ |

The optimized parameters were used for multiplexed imaging of protein targets in *E. coli* and effector mRNAs in the pathogen *Y. enterocolitica*. The 4-plex Exchange-PAINT imaging enabled us to visualize MreB and the RNA polymerase β’ subunit RpoC in correlation with the cell envelope (LPS) and the nucleoid (Figure 3A), capturing several features of subcellular organization. First, it demonstrates the correlation of RpoC and the nucleoid in 3D space [42,59], as revealed by a positive CBC value of 0.62 ± 0.08 (Figures 3 and S8). Similarly, multiplexed imaging captured the spatial proximity of MreB and LPS (CBC = 0.3 ± 0.2), which are localized to the inner and outer membrane, respectively (Figures 3 and S8). Here, the high 3D resolution of DNA-PAINT clearly visualizes the separation of inner and outer membrane, reflected by a peak-to-peak distance of 51 ± 12 nm of both signals (Figure 3E). Taking labeling accuracy, localization precision and the positioning of MreB and LPS on opposite sides of inner and outer membrane into consideration, this value leads to an estimated periplasm size of ∼ 20 – 30 nm, well in agreement with electron microscopy studies [51]. We recently reported nucleoid positioning close to the membrane in fast-grown *E. coli* cells [48]. 3D averaging of cross-sections of the cell cylinder revealed the ring-like nucleoid distribution with low signal at the radial cell center observed before. Here, we determined a distance of 129 ± 31 nm between LPS and DNA signal, being in good agreement with the measurements obtained in single-slice 2D imaging [48].

Having optimized DNA-PAINT imaging on rRNA-FISH, we transferred the conditions to multiplexed 3D imaging of effector mRNA targets in *Y. enterocolitica*. We wondered whether these transcripts are spatially organized, in particular whether they are localized towards the membrane. Such a positioning might mediate direct insertion of nascent effector proteins into the secretion machinery, thus streamlining effector protein secretion. In agreement with our previous work, we did not find evidence for such an organization for *yopM*. The number of clusters, which we take as a proxy for transcript abundance, amounted to 40 ± 17, in good agreement with the value we previously reported for this growth condition [25]. Extending this work to the mRNAs of effector proteins YopE and YopH, we did not find any evidence for transcript membrane localization either, but roughly two times more *yopE* and *yopH* clusters compared to *yopM* (Figure 4C). These values are in good agreement with the relative abundance of these effectors in the bacterial proteome [60] and secretome [61]. Furthermore, the transcripts show a slight spatial correlation (CBC values of 0.15 for *yopE*-*yopH*, 0.24 for *yopH*-*yopM* and 0.10 for *yopE*-*yopM*) and a preferentially cytoplasmic localization towards the cell poles (Figure 4E and F, Supplementary Video 3), in agreement with separation from the center-localized nucleoid in secreting cells observed in recent work [25].

In summary, we report an easy-to-establish platform and straightforward approach for optimizing DNA-PAINT imaging in bacterial cells. Our screen revealed docking-imager strand pairs that allow 3D DNA-PAINT imaging at 15 - 20 nm resolution, even though we were limited to the use of the suboptimal DNA-PAINT dye ATTO655 [38]. Exchange-PAINT imaging in *E. coli* and *Y. enterocolitica* showcase that multiplexed 3D super-resolution imaging provides meaningful biological insights that are otherwise hidden by the diffraction limit of light.

In the future, our platform can easily be used to screen novel sequences, or to test different dyes on systems that provide optical sectioning [38]. The use of novel DNA-PAINT strategies, such as fluorogenic probes or left-handed DNA [16,17], as well as artificial intelligence can further improve resolution, acquisition speed and multiplexing capability [62–64]. We hope that our work motivates bacteriologists to make use of the high resolution and multiplexing capabilities provided by DNA-PAINT, and that this gives rise to important biological insights.

## Methods

### Bacterial cell culture

The bacterial strains used in this study are listed in Table S1. *E. coli* strains were grown in LB Lennox (Carl Roth, X964.2) at 37 °C with shaking at 200 rpm. Cloning and conjugative plasmid transfer used donor strain ST18, which is auxotrophic for 5-aminolevulinic acid (ALA) [65]. Cells were grown in LB medium at 37°C with shaking at 200rpm. ST18 was supplemented with 50 µg/mL ALA. Kanamycin was used at 50 µg/mL. Overnight cultures were inoculated from glycerol stocks, diluted 1:100 on the next day into fresh LB and grown to mid-exponential phase (OD_600_ reached ∼0.7) before chemical fixation.

*Y. enterocolitica* strains were grown and expression of the T3SS was performed as previously described [66] with the following modifications and additions: The initial optical density at 600 nm (OD_600_) of the cultures was set to 0.15 for secreting and 0.12 for non-secreting conditions; bacteria were incubated at 28° for 90 min before being incubated at 37° for further 150 min to induce expression of the *yop* regulon (structural components of the T3SS).

### Plasmid construction

Chromosomal modifications were introduced by two-step allelic exchange [67] with suicide plasmids based on the vector pEB17 (pDS132 backbone) [68]. All plasmids were assembled by Gibson/HiFi assembly. Plasmids used in this study are listed in Table S1 and primers in Table S2.

pMBP-36 (RpoC-ALFA): The pEB17 backbone was linearised by PCR with primers KA-21/KA-22 using Q5 High-Fidelity DNA Polymerase (NEB, M0491L). The template was removed by DpnI digestion (1 h, 37 °C, followed by 20 min heat inactivation at 80 °C; NEB, R0176L). The upstream and downstream homology arms flanking the insertion site in *rpoC* were amplified from MG1655 genomic DNA with primers KA-3/KA-17 and KA-8/KA-18, respectively. The ALFA-tag [50] coding sequence was generated by annealing two complementary oligonucleotides (1 µM each in 50 mM NaCl; 95 °C for 5 min, then successively cooling to 23°C at increments of 1 °C/min for 72 cycles, then keep at 4 °C).

pMBP-58 (MreB-sfGFP): The pEB17 backbone was amplified with KA-21/KA-22. The insert, comprising the *mreB* region flanking the insertion plus the sfGFP cassette (MreB-sfGFP-MreBCD) was amplified with RX-36/RX-37 from genomic DNA of *E. coli* NO34 [69]. The template was removed by DpnI digestion (1 h, 37 °C, followed by 20 min heat inactivation at 80 °C).

All Fragments were purified for following Gibson assembly using commercial PCR cleanup kits (Macherey-Nagel, 740609). Fragments were combined with NEBuilder HiFi DNA Assembly Master Mix (NEB, E2621L). The assembly was transformed into ST18 using electroporation. Colonies were screened by colony PCR and plasmids from positive clones were isolated and verified by sequencing.

### Chromosomal integration by conjugation and allelic exchange

Chromosomal modifications were introduced by two-step allelic exchange [67]. Plasmids were transferred from the ALA-auxotrophic donor strain ST18 into the recipient strain MG1655 by conjugation. Donor and recipient cultures were grown to mid-exponential phase (OD₆₀₀ ≈ 0.5-0.8). Donor cells were washed three times in LB to remove residual ALA and mixed with recipient cells at a 3:1 OD_600_-normalised ratio. The mixture was resuspended in 300 µL LB, spotted on LB agar without ALA, and incubated overnight at 37 °C. Cells were then resuspended in 1 mL LB and plated on LB containing 50 µg/mL kanamycin to select for plasmid integration. Single colonies were grown overnight in LB without antibiotics to allow the second recombination, then plated on LB containing 10 % sucrose and incubated at 30 °C to select against cells retaining the *sacB*-containing plasmid backbone. Sucrose-tolerant colonies were restreaked on LB + sucrose and LB + kanamycin, and kanamycin-sensitive clones were screened by colony PCR with primers flanking the modified locus (KA-64/KA-49 for *rpoC*-ALFA; RX-47/RX-48 for *mreB*-sfGFP).

### Cell preparation

The RNA-FISH protocol was adapted from Skinner et al. [33]. For fixation, 6 OD_600_ units of cells in exponential phase were mixed 1:10 with 37% Formaldehyde (FA, Sigma-Aldrich, 252549) to obtain a final concentration of 3.7% and incubated for 30 minutes at room temperature on a nutator. Cells were harvested by centrifugation (400x g, 8 min); the supernatant was discarded, and the cells resuspended in 1mL RNase-free PBS (Thermo Fisher Scientific, AM9625). The cells were washed twice with 1mL of PBS again. After the last wash, the cells were resuspended in 300µL of RNase-free ddH_2_O (Sigma-Aldrich, 9610) and 700µL RNase-free Ethanol (Sigma-Aldrich, 51976-F) was added to permeabilize the cells. Samples were incubated for 1 h at room temperature on a nutator and subsequently used directly for immobilization or stored at 4°C until further use.

### Immobilization and hybridization

For immobilization of cells, 8-well chamber slides (Sarstedt, 94.6190.802) were etched for one hour with 3M KOH and then washed thoroughly with ddH_2_O. Slides were incubated with freshly prepared 0.01% poly-L-lysine (Sigma-Aldrich, P8920, 1:10 dilution) for 15 minutes, washed with ddH_2_O, air-dried in a dust-free environment and finally UV-sterilized. All subsequent steps were performed with RNase-free solutions.

50 µl of permeabilized cells in solution were washed once with PBS, taken up in 400 µl PBS, added to PLL-coated chamber slides and incubated for 30 minutes at room temperature with slow nutation. Afterwards, cells were removed and the chambers were washed twice with PBS. Chambers were then incubated without shaking for 30 min with a 1:100 solution of UV-sterilized gold nano-urchins (Cytodiagnostics, GU-90-20) in PBS. The unsettled nano-urchins were removed by washing the chambers once with PBS. Samples were then blocked for 10 minutes using 1% RNase-free BSA (Carl Roth, 3737.2) in PBS and subsequently washed twice again with PBS.

For hybridization with the FISH-probes, chambers were washed once with 40% washing buffer (for 1mL: 353µL 100% Formamide (Carl Roth, P040.1), 100µL 20x SSC (Thermo Fisher Scientific, AM9763), 547µL RNase-free ddH_2_O), followed by an incubation of at least 10 minutes in 40% washing buffer. Afterwards, the hybridization solution containing FISH probes was incubated for 16 hours at 30°C in a sealed petri dish. All RNA-FISH probes used in this study with the corresponding concentration are listed in Table S2. On the next day, the hybridization solution was removed and the chambers washed twice with fresh 40% washing buffer at 30°C for 30 minutes each. Finally, samples were washed twice with PBS and stored at 4°C until imaging.

### Preparation of hybridisation solution

The hybridization buffer was prepared as a 10 ml batch according to Skinner et al. [33]. Dextran sulfate (1 g; Sigma-Aldrich, D8906) was dissolved in 5 mL DEPC-treated water in a 50 mL polypropylene conical tube by vortexing, and the solution was mixed on an orbital shaker for ∼30 min until bubbles had cleared. Frozen aliquots of formamide were equilibrated to room temperature before opening. Formamide (3,530 µl), salmon sperm DNA (10 mg; Sigma-Aldrich, 31149-F), 20× SSC (1 ml), 200 mM VRC (100 µl; Sigma-Aldrich, 94742) and BSA (40 µl of a 50 mg/ml stock) were then added, and the volume was adjusted to 10 ml with DEPC-treated water. The buffer was nutated until it was mixed homogeneously, sterilized through a 0.22 µm filter, and stored at −20 °C for up to one year. Final concentrations were 10% (w/v) dextran sulfate, 35% (v/v) formamide, 1 mg/ml salmon sperm DNA, 2× SSC, 2 mM VRC and 0.2 mg/ml BSA.

### Nanobody staining

Bacteria were fixed in mid-exponential phase by directly adding fixation solution, resulting in a final concentration of 33mM NaPO4 (pH 7.5), 2% methanol-free FA (ThermoFisher, 28908) and 0.1% Glutaraldehyde (GA, Carl Roth, cat-4157.2). Solutions were incubated for 15 min at RT. Bacteria were then pelleted by centrifugation (3 min, 5,000x g), washed once with PBS and quenched for 20 min using 50mM NH_4_Cl in PBS. Bacteria were immobilized on APTES-coated and GA-activated chamber-slides, which were prepared as following: Chamber slides were etched 1 h at RT with 3 M KOH, washed 3x with ddH_2_O and coated for 1 min with freshly prepared 2% APTES in ddH_2_O. Chamber slides were washed twice with ddH_2_O and cured overnight at 60°C. On the next day, surfaces were activated by incubating for 15 min with 1% glutaraldehyde in ddH_2_O. Chambers were washed twice with ddH_2_O and excess GA was quenched for 20 min with 50mM NH_4_Cl in PBS, blocked and permeabilized with 3% BSA and 0.5% Triton-X100 for 1 h at RT. Nanobody staining was carried out in 1% BSA and 0.1% Triton-X100 overnight at 4°C. Nanobodies with an ectopic single cysteine were purchased from NanoTag (N0305-250UG for @GFP clone 1H1, C8000 for @GFP clone 1B2 and N1505-250UG for @ALFA nanobodies) and conjugated as described by Sograte-Idrissi *et al*. [70]. To label ALFA-tagged RpoC and MreB^sw^-sfGFP, nanobodies carrying the respective docking strands were used in a concentration of 25 nM. LPS was stained using 10 µg/ml rabbit-anti-*E. coli* primary antibody (Bio-Rad, 4329-4906) functionalized with DNA-PAINT docking strands via Oligo FlexLinker 2.0 kit (ProteinTech, KFA151). FlexLinkers were functionalized with DBCO-modified oligos (metabion) according to the manufacturer’s instructions. After incubation with nanobodies or antibodies, the cells were washed twice with staining buffer, post-fixed with 2% MeOH-free FA, washed with PBS and quenched again for 20 min with 50mM NH_4_Cl in PBS. For drift-correction, a dilution of 1:100 of gold nano-urchins was added to the slides as described above.

### DNA-PAINT imaging

DNA-PAINT images were acquired on a Vutara VXL 3D super-resolution microscope equipped with 405 nm, 488 nm, 555 nm and 640 nm excitation lasers, a Hamamatsu Fusion BT scientific complementary metal–oxide–semiconductor (sCMOS) detector, a 60× silicon oil objective (NA 1.3) and biplane optics. The system was operated at 26°C. SRX analysis software (7.16) was used for single-molecule data analysis. MLE fitting using a model generated with immobilized fluorescent fiducial markers (100 nm TetraSpecks, ThermoFisher, T7279) was used for localization with thresholds varying with docking-imager strand sequences.

For comparison of all imager strands, DNA-PAINT images were acquired for 20,000 frames at a laser density of 8.3 kW/cm² (640 nm) and an exposure time of 50 ms. The imager strands conjugated with ATTO655 were diluted to the corresponding concentration in imaging buffer (500mM NaCl, 1mM EDTA, 0.02% Tween-20, in PBS). The samples were equilibrated with the imager buffer for two to three minutes before adding fresh solution containing the DNA strands.

In Exchange-PAINT experiments, chambers were washed carefully with imaging buffer on stage after every acquisition (5 times or until no residual blinking was observed) before addition of the new imager strand. Brightfield, GFP and Cy3 reference images were recorded before DNA-PAINT acquisition.

### Automatic image analysis pipeline

Data analysis was carried out using a self-developed, fully automated and batch-capable Python pipeline, parts of the analysis are based on the Picasso v11.0 framework [5]. Localisations were drift-corrected using the adaptive intersection maximization (AIM) algorithm [71] as implemented in the Picasso package [5]. Bacterial cells were segmented from brightfield images using Cellpose-SAM (cpsam model, Cellpose 4.0.9; nominal diameter 20 px, flow threshold 0.4, cell-probability threshold 0.0) [72]. To register DNA-PAINT localizations to segmentation masks, a global six-parameter affine transform was fitted by optimizing the overlap of the localization cloud with the binary segmentation masks [73]. The calculated transform matrix was reused for all downstream channel registrations, which ensured consistent registration and mitigated chromatic aberration and other offsets between optical paths. The Cy3 reference image was warped into mask space using the inverse of this transform.

Cells unsuitable for quantification were removed sequentially, including:

- Cells close to the image borders or cut cells (bounding box closer than 5 px to the image borders).
- Cells containing gold fiducials (median photon count exceeded 5× the median photon count of the entire field of view).
- Mechanically unstable cells, that result in a motion-blurred super-resolution image. Per-cell localizations were split into ten time-bins of similar size, and the per-cell centroid displacement between bins was calculated. Cells with displacements > 3 SD, calculated from all cells in the field of view, were excluded.

### Calculation of experimental localization precision (σ_NeNA_)

σ_NeNA_ was calculated globally and per cell using the function implemented in Picasso [36]. Subsequent-frame localizations were linked into individual binding events using the Picasso linking module with a maximum dark time of 3 frames and a linking radius of 130 nm. Per-cell metrics automatically extracted by the custom pipeline include localization count, median localization precision and photon count, number of binding events, event duration in frames as well as localizations per event.

### Correlation of DNA-PAINT signals with FISH staining

For normalization, the Cy3 image was background-corrected by subtracting the median of all non-cell pixels (negative values clipped to zero), and the integrated intensity within each cell mask was measured. Per-cell localization count (or binding-event count after linking) were normalized by this value. Cells with negative intensity after background-correction were removed from the dataset (mostly relevant for negative controls). As no absolute Cy3 calibration was performed, these ratios are relative measures of binding activity, valid for comparison within consistently acquired datasets. Cell morphology and intensity metrics (area, bounding box, centroid, integrated Cy3 intensity) were extracted from the label mask using regionprops from the scikit-image library [74]. Localization-derived per-cell statistics were computed and aggregated using the Python libraries NumPy [75] and Pandas [76].

### Determination of DNA-PAINT hybridization kinetics

DNA-PAINT image sequences were analyzed using the automatic image-analysis pipeline described earlier. The ON-times of all binding events were extracted from the localization lists and events lasting only a single frame were filtered. The ON-time of each event duration was converted to milliseconds by multiplication with each acquisition’s frame exposure time.

For each condition, ON-time distribution fitted with a two-population (biexponential) decay model (equation 1):

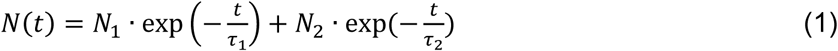

Here, *τ*_1_ and *τ*_2_ are exponential decay time constants. For the slow component, the apparent dissociation rate is *k*off = 1/*τ*, and the corresponding binding half-time is *t*_1/2_ = *τ*_slow_ ln2. N_1_ and N_2_ represent the corresponding amplitudes. The fast component (*τ*_fast_) was attributed to short-lived, nonspecific binding events, and the slow component (*τ*_slow_) to specific hybridization events. This two-population assignment was validated against unstained (no complementary target) control samples. ON-time distributions of the control experiments were fit with a mono-exponential decay model as the primary model. The resulting half-time was compared with the fast component of the stained samples’ bi-exponential fit.

Binding-event kinetics were computed both as a single pooled fit per condition (all events from all cells combined) and per individual cell, to obtain a cell-to-cell distribution of dissociation rates rather than one value per condition. For the per-cell analysis, the bi-exponential fit above was repeated independently on each cell’s own retained events; cells with fewer than 300 retained events were excluded, as the bi-exponential fit becomes unstable below this count. The ON-time, τ_B_, was taken as the slow component of each cell’s fit (specific binding), and the per-cell dissociation rate constant computed as established in DNA-PAINT imaging [5,20] (equation 2):

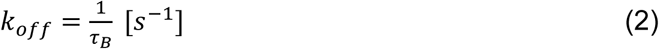

Values were reported as the mean ± SD of the per-cell *k*_off_ distribution (capturing genuine cell-to-cell variability).

### Coordinate based colocalization

Localizations were first filtered by segmentation masks. For each acquisition, channels (DNA-PAINT, BF and Cy3) were registered using Picasso’s redundant cross-correlation algorithm applied to the entire field of view [77]. CBC scores were computed on a per-cell basis and in 2D. For each cell, we extracted the localizations from the 200 nm center slice (±100 nm of each channel’s median z-value) for all aligned channels. The coordinate-based colocalization algorithm from Malkusch *et al*. was then applied [47]. Cells with fewer than 20 localizations in either channel were excluded from analysis. Positive controls were obtained by splitting the events by odd/even frame parity, and computing CBC between the two datasets. All analysis was implemented in Python (NumPy/SciPy cKDTree for spatial queries, Picasso 0.11 for registration and event linking).

### Line-profile and cross section analysis

For each cell, the segmentation mask was skeletonized and the longest end-to-end path was taken as the long axis. A smoothing spline was fitted to this path, and 10 % of its length was trimmed at each pole to exclude the hemispherical caps. Each localization was assigned to its nearest point on the spline in the image plane, and its signed transverse distance from the axis was calculated perpendicular to the local spline direction. The four channels were registered to the LPS channel with Picasso (RCC); a residual rigid translation was corrected by aligning per-cell localization centroids to the LPS channel. Only localizations with an axial precision better than 35 nm and a lateral precision better than 15 nm were used.

For line profiles, the distribution of signed transverse distances of each cell was estimated by a Gaussian kernel density estimate (bandwidth 10 nm), so that each cell contributed equally regardless of its number of localizations. The per-cell densities were averaged across cells (N_cells_ = 14) and are shown as mean ± SD., normalized so that the mean curve of each channel peaks at 1.

For cross-section averages, the axial position (z) of each localization was recentered on the median z of its cell. Localizations from all cells (N_cells_ = 14) were pooled per channel, binned in 2D (10 nm bins) as a function of signed transverse distance and z, smoothed with a Gaussian kernel (σ = 10 nm), and normalized to a maximum of 1.

### Measuring distances between DNA-PAINT channels

Distances measurements for Exchange-PAINT images in *E. coli* were performed using Fiji (ImageJ) [78] using a custom macro. Because DNA-PAINT LPS labeling is discontinuous around the cell envelope rather than a continuous line, cell outlines were not extracted by direct thresholding of the full outline. Instead, a small Gaussian blur (σ = 1.5 px) was applied to the LPS channel, followed by thresholding using the Otsu algorithm [79], and individual labeling clusters below a minimum area (8 px²) were discarded as noise. For rod-shaped, non-dividing cells, all retained clusters were pooled and enclosed within their two-dimensional convex hull. Dividing cells (with a septum) and cells with pronounced curvature, whose outline was not globally convex were manually subdivided into two or more segments along their length; each segment’s LPS clusters were enclosed by their own convex hull, and the union of the segment hulls was taken as the final outline.

We traced the outer boundary of each cell’s LPS signal and smoothed it with a spline. Because this boundary lies at the outer edge of the signal, we shifted it inward by 6 pixels, so that the line runs through the middle of the LPS signal instead of along its edge.

LPS and MreB signals were straightened using the outlines generated above and using a line width of 100 px using the “Straighten” function in Fiji. For each channel, the straightened image was averaged along its perimeter axis to yield a single one-dimensional radial intensity profile (mean intensity as a function of radial distance from the outline). For each cell, the periplasmic gap was calculated as the absolute difference between the LPS and MreB peak positions, converted to nanometers using the image’s pixel information. Radial intensity profiles and peak positions were exported per cell for independent verification. LPS/DNA distances were calculated similarly, but using a 1/e threshold as published previously [48].

### DBSCAN analysis and counting of mRNA clusters

For DBSCAN analysis in *Y. enterocolitica* RNA-FISH experiments, localization files were preprocessed as described above. Data was visualized using Picasso v0.11 [5]. The DBSCAN algorithm requires to set two input parameters, the searching radius ε as well as min. points, the minimum number of localizations that form a cluster [56,80]. To account for anisotropic localization precision, we used an ellipsoidal neighborhood defined by separate lateral and axial distance thresholds for 3D DBSCAN analysis (ε_xy_ and ε_z_). These values were determined based on the measurements’ σ_NeNA_ value (2x σ_NeNA_, ∼13nm, ε_xy_), as well as on the axial localization precision σ_z_ provided by SRX (2x σ_z_, ∼ 50nm, ε_z_). To experimentally determine min. points, ∼300 apparent mRNA clusters per species were manually selected using the Picasso “Pick” tool on linked localization data. The median number of points per cluster was chosen as min points (7 for yopM, 14 for yopH, 16 for yopE). DBSCAN analysis was then carried out on all datasets using these parameters.

### Generation of 3D cluster maps

To relate cluster position to cell geometry, the DNA-PAINT image of the EUB338 channel served as a whole-cell reference and was used to fit each cell as a three-dimensional sphero-cylinder. The cell center axis was determined from the reference-channel localizations, and cell boundaries were approximated by a sphero-cylinder enclosing 95% of localizations radially and 99% axially, dilated by 50 nm into each direction to include all localizations belonging to the cell. As DNA-PAINT channels were recorded in separate rounds, localization files were slightly misaligned. We estimated this shift as the median centroid displacement between channels across matched cells and corrected the reference channel accordingly.

Within each cell, cluster positions were described relative to a fitted rod-shaped outline using the position along the long axis together with the distance and the angle from the axis. Positions were scaled to each cell’s dimensions, which allowed us to overlay clusters from all cells within a cell model with cell dimensions representing mean values extracted from the single-cell datasets [56,81].

### Use of large-language models (LLMs)

LLMs were employed to assist with several tasks. Claude Opus (versions 4.5 – 5.5) was used to assist with coding. All generated or modified code was rigorously checked, and results were inspected for potential artifacts. LLMs were further used to assist in writing.

## Data and code availability

At this point, all data and code used in this study is available from the authors upon request. Data and code will be openly shared in the next version of this manuscript.

## Acknowledgements and funding

The authors thank Francesca Ermoli and Corentin Brianceau (Max Planck Institute for Terrestrial Microbiology, Karlsruhe Institute of Technology) for help with initial sample preparation optimization. K.A. and C.S thank Til Schipniewski for help with sample preparation and the entire Zimmer group for the great atmosphere and constant support. K.A., R.X. and C.S. acknowledge funding by the State of Bavaria via the Distinguished Professorship Program (DPP, 3.1-3122.05-008/014 P000122427). C.S. further acknowledges funding by the University Würzburg via a JMU Seed Grant. A.D. acknowledges funding by the German Research Foundation (DFG, grant DI 1765/5-1, project number 456615895).

## Author contributions

C.S. and K.A. conceptualized the study. K.A. performed the majority of data acquisition and analysis. P.K. performed and analyzed the initial RNA-FISH experiments. R.X. and K.A. generated bacterial strains. A.S., K.A., and P.K. prepared samples. C.S. and A.D. acquired funding. K.A. and C.S. wrote the manuscript with input from all co-authors.

