## Supplementary Information for "A simple platform for *in situ* DNA-PAINT optimization enables improved multiplexed super-resolution imaging in Gram-negative bacteria"

Affiliations:

Figures S1 – S11

Tables S1 – S2

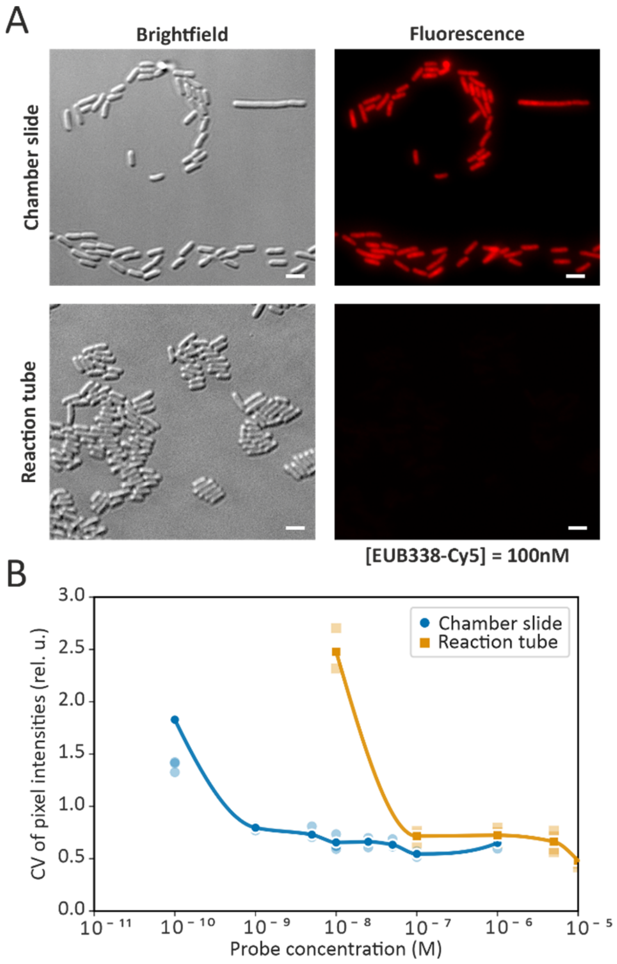


***Figure S1: Optimization of RNA-FISH staining procedure****. (A) exemplary images of E. coli stained with 100nM EUB338-Cy5 in chamber slides (top row) and in reaction tubes (bottom row). (B) Comparison of heterogeneity of staining by coefficient of variation (CV) of cells stained on chamber slides (orange) and in reaction tubes (blue) (Mean CV + SD calculated between individual cells for each condition). Scale bars are 5 µm.*

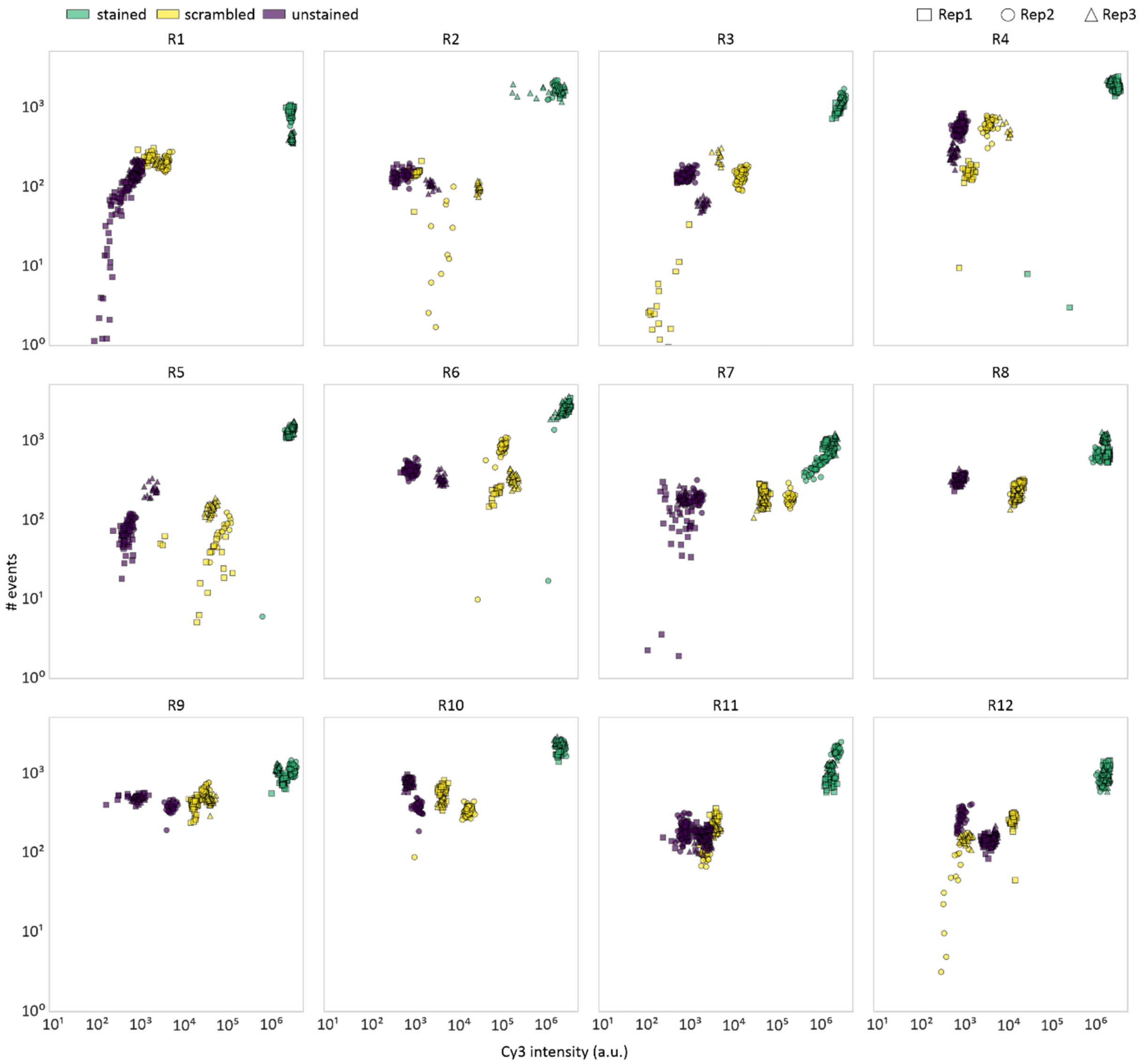


***Figure S2: Correlation of DNA-PAINT signal with Cy3 intensity in EUB338-stained E. coli cells****. Correlated are the number of events against the Cy3 intensity for each cell under stained, scrambled and unstained conditions. $N_{replicates}$ = 3.*

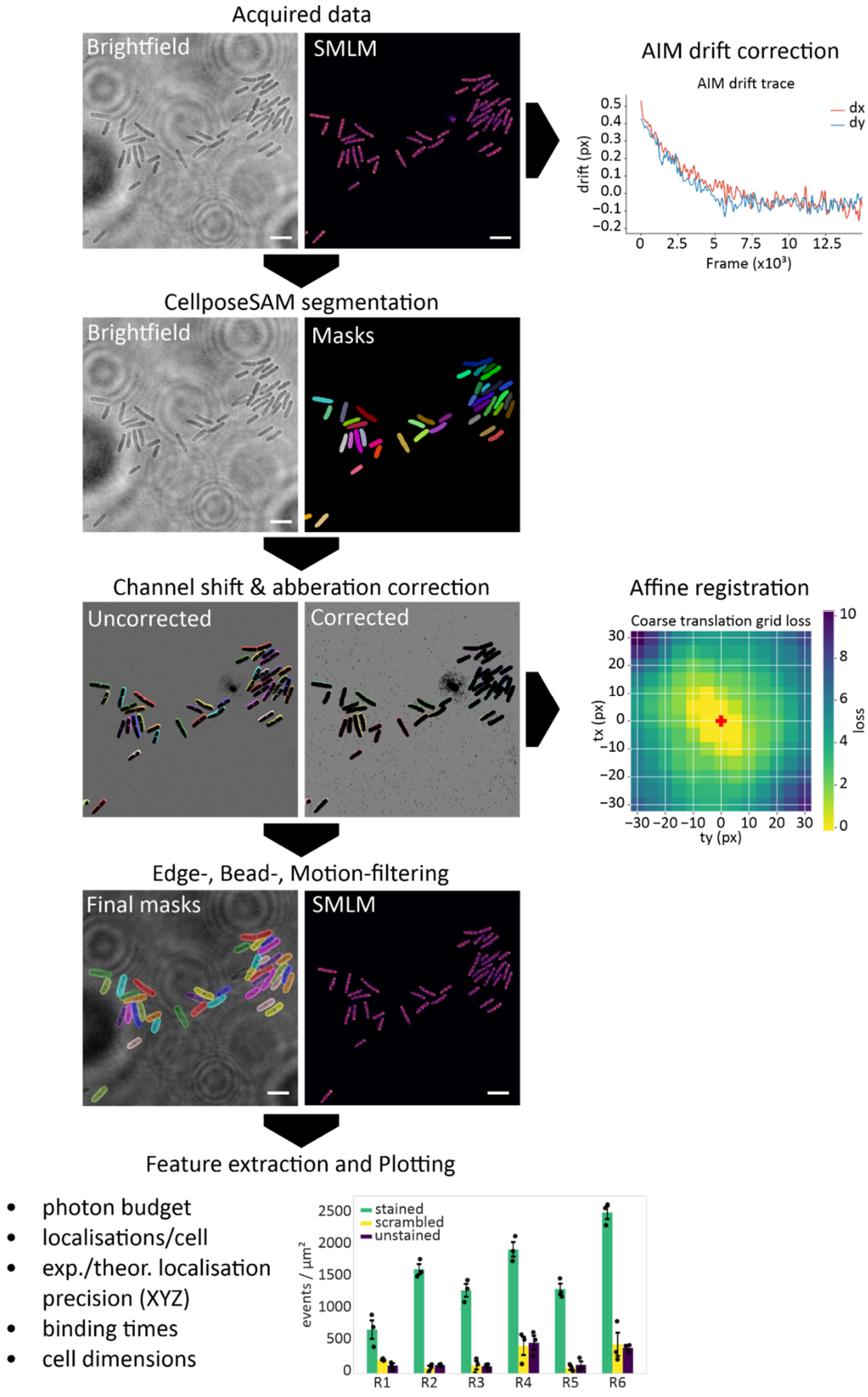


***Figure S3: Schematic workflow of image-analysis pipeline**. Localization lists of DNA-PAINT measurements are firstly drift-corrected by the AIM-algorithm [1]. The corresponding brightfield images are segmented using CellposeSAM to generate binary masks. To correct for channel shifts and chromatic aberration, single-molecule localizations are registered with the segmentation masks using an affine transformation matrix. Cells are filtered for position (excluding cells closer than 5 px to the border), attached fiducial markers / beads (excluding cells with > 5x median photon count), and motion (excluding cells with centroids moving > 3x SD over the acquisition). Single-cell data is extracted from remaining masks, followed by NeNA analysis and linking of subsequent-frame localizations. Parameters then get extracted from the linked and filtered localization files and plotted. Scale bars are 5 µm.*

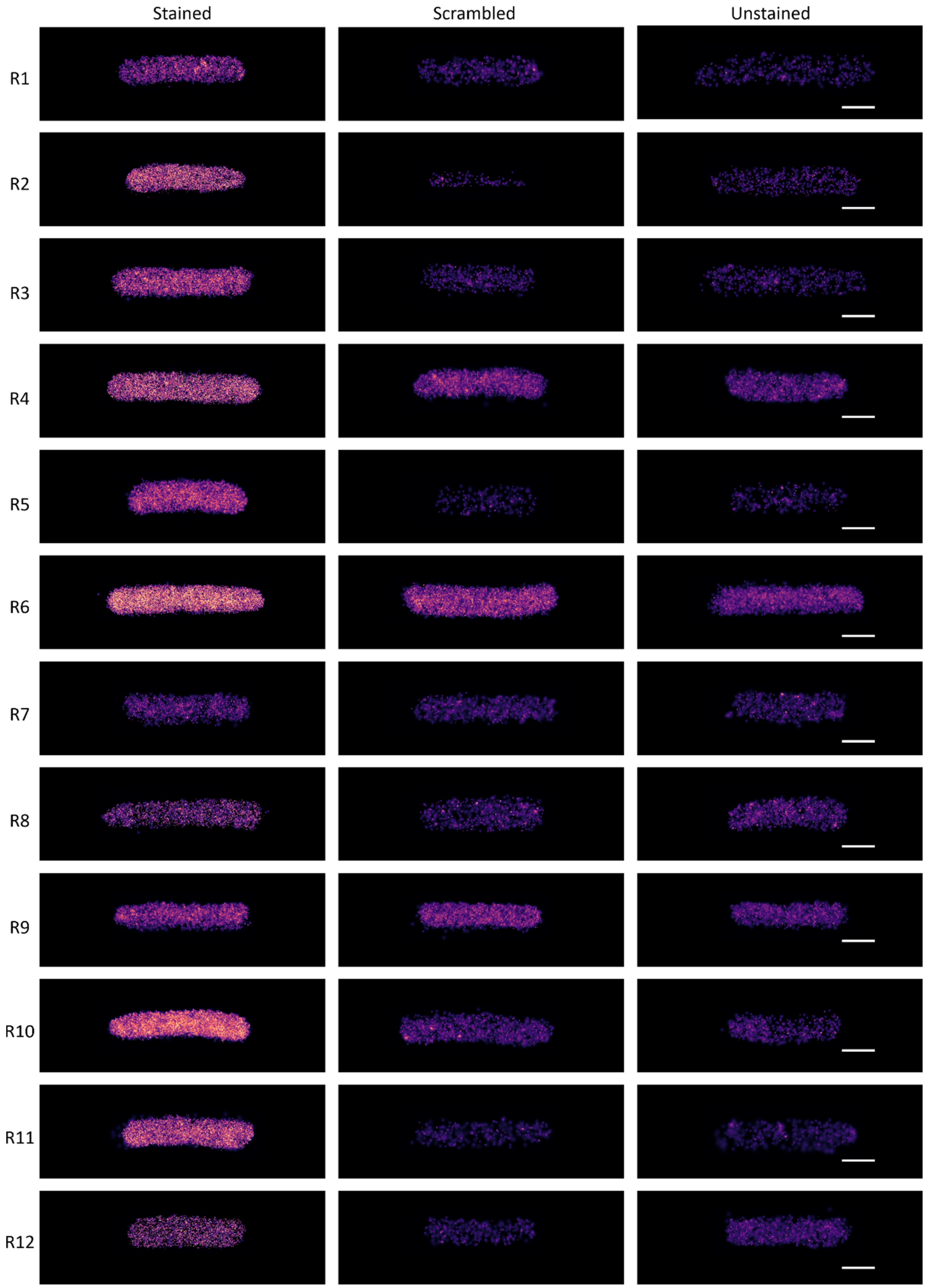


***Figure S4: Representative DNA-PAINT reconstructions for all tested imager-strands R1-R12 and under the different staining conditions (stained, scrambled, unstained)****. Images were rendered in Picasso using the same contrast settings and by isotropic localisation precision. Scalebars are 1µm.*

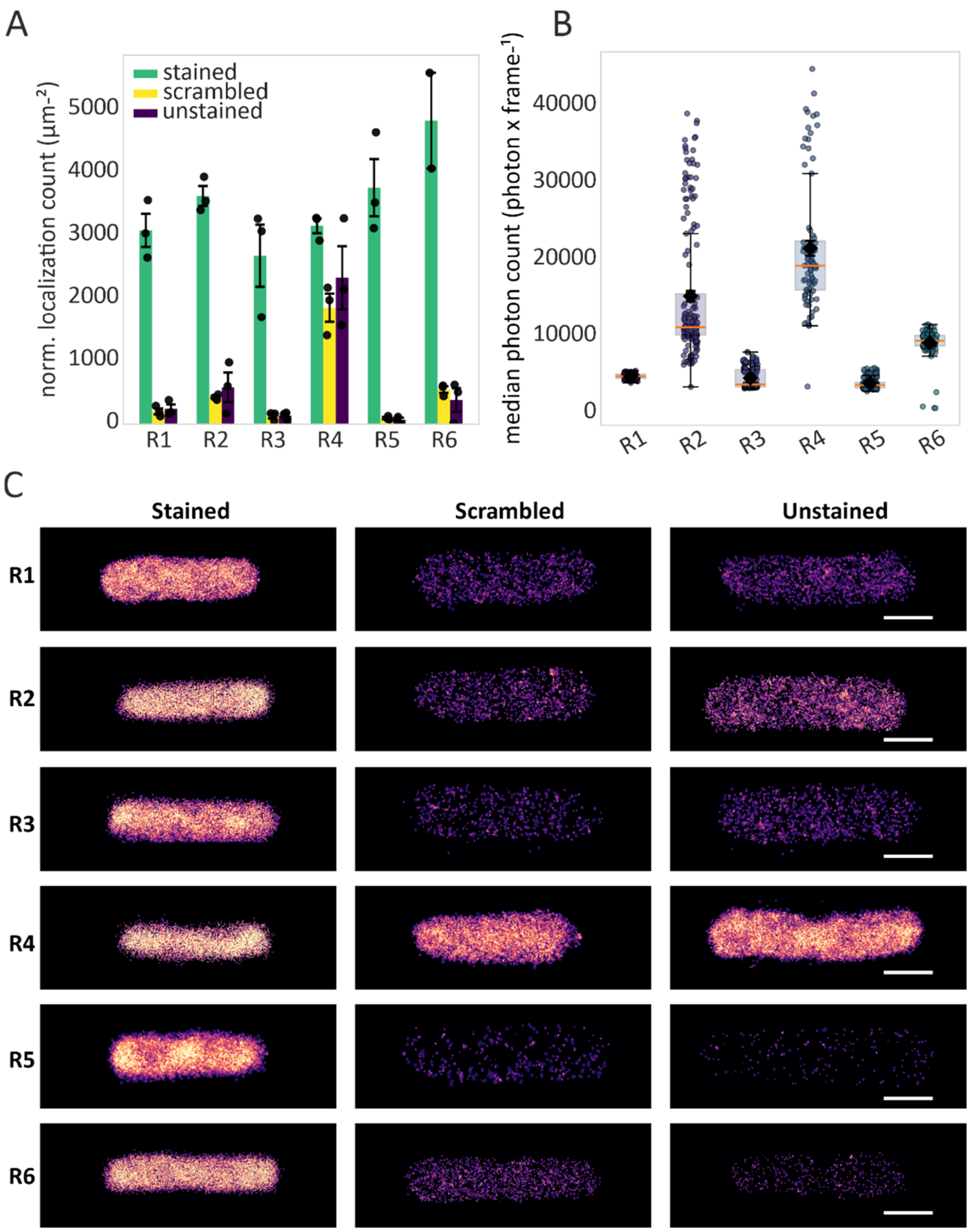


***Figure S5: Assessing imager strand performance using 1nM imager strand concentration****. (A) Comparison of the normalized localization count (events/µm²) for each imager strand. Area normalization was performed using the area of segmentation masks. (B) Comparison of single-molecule signal brightness for imager strands R1 – R6. Note that the brightness for R2 and R4 are affected by overlapping single-molecule signals, and thus do not reflect the actual signal brightness. Accordingly, the number of binding events for R2/R4 in (A) are also underestimates. (C) Representative DNA-PAINT reconstructions of individual cells. Images were rendered in Picasso using the same contrast settings and by isotropic localisation precision.* $N_{replicates}$ *= 3. Scalebars are 1µm.*

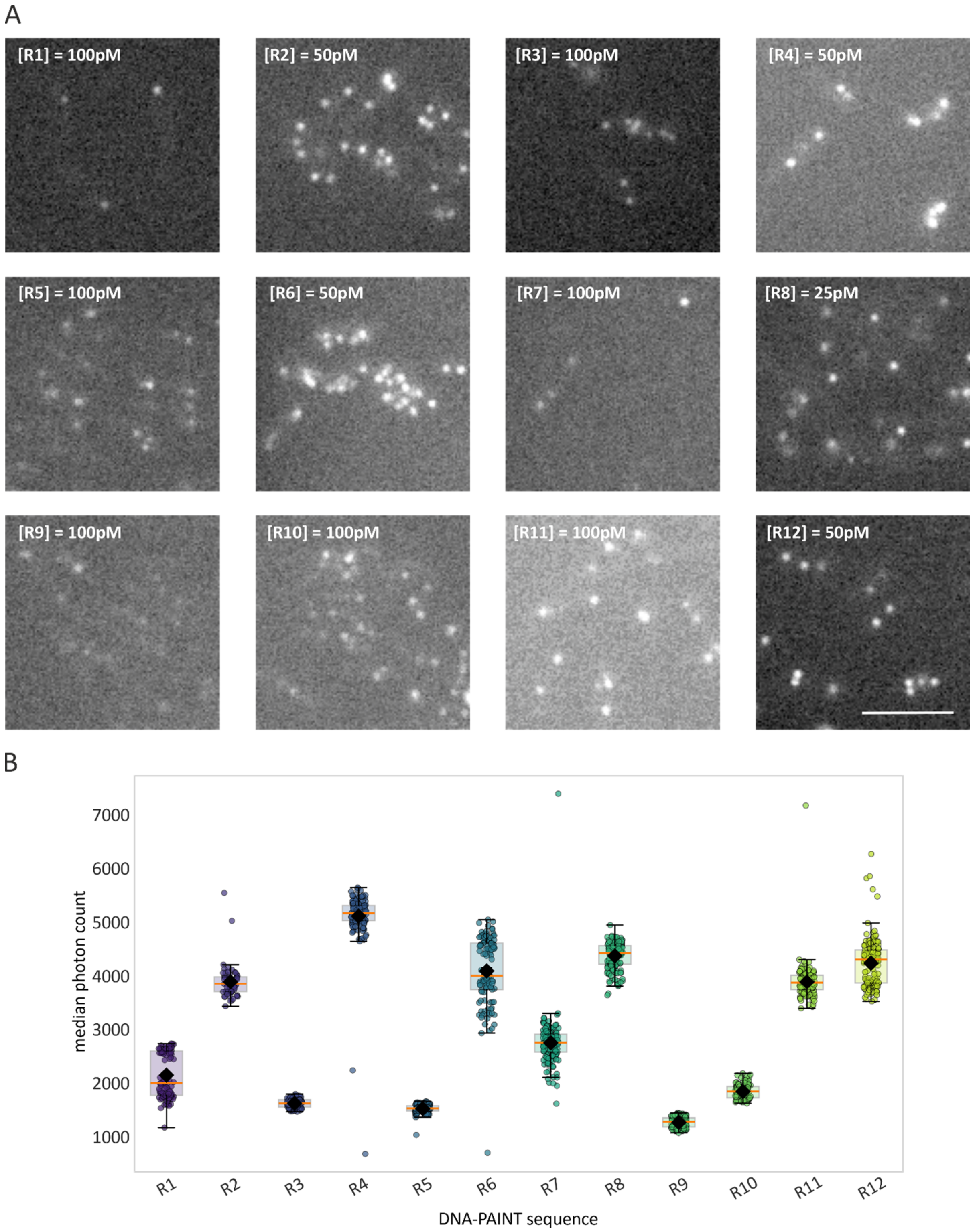


***Figure S6: Brightness varies with DNA-PAINT sequence**. (A) Single frames of DNA-PAINT measurements recorded with the indicated imager strand concentrations and 50 ms exposure time. (B) Median photon count per frame for each imager strand. Data points represent single cells, the orange line represents the median, boxes the 25 and 75% percentiles and whiskers show the 99% percentile. $N_{replicates}$ = 3. Scale bar in (A) is 5 µm.*

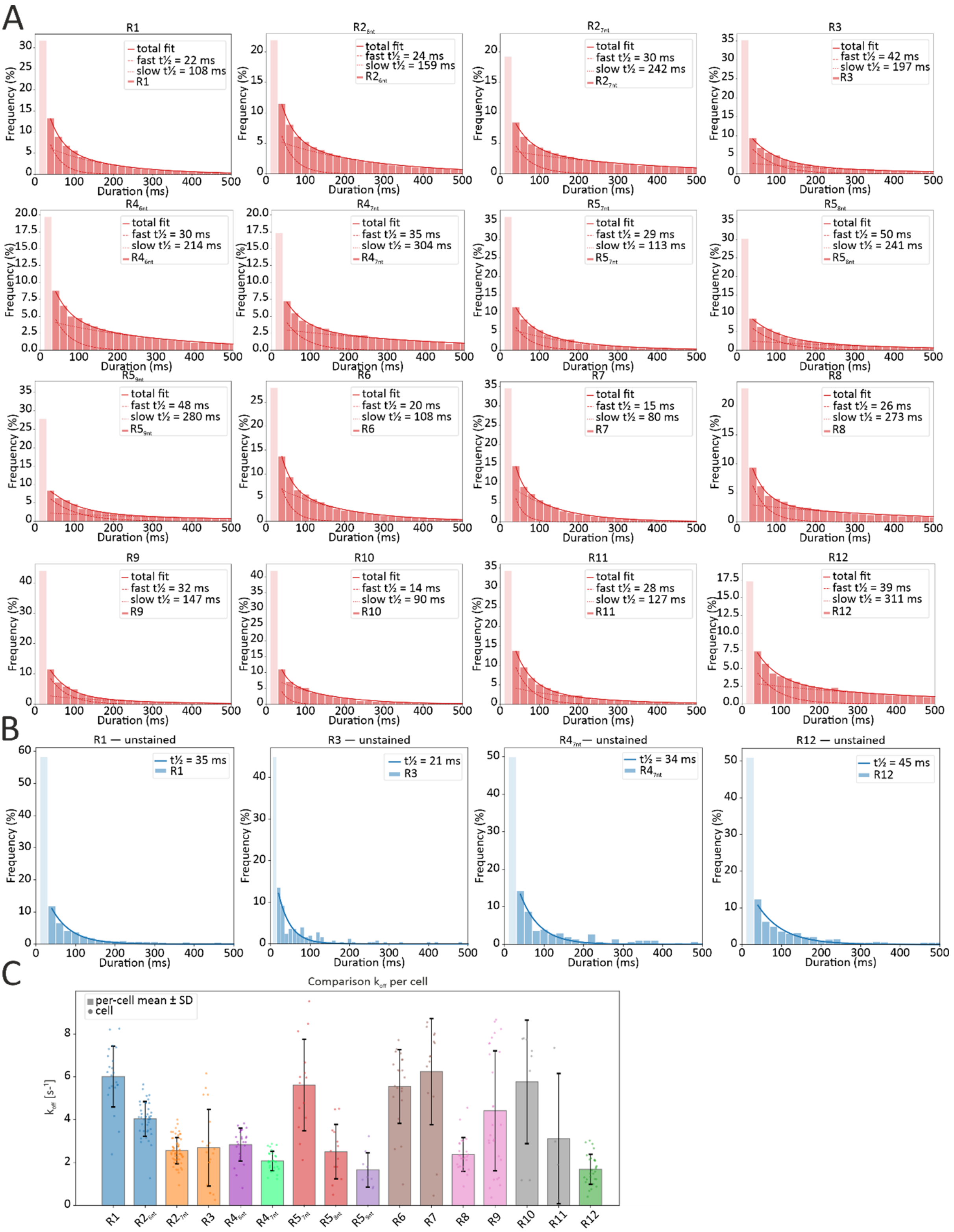


***Figure S7: ON-time analysis for the DNA-PAINT sequences R1-R12**. (A) ON-time distribution for the indicated DNA-PAINT sequences. Data was fitted with a bi-exponential decay function. Slow and fast species of binding half times $\tau_{1/2}$ represent specific and unspecific binding of imager strands, respectively. (B) ON-time distribution for unstained controls of selected imager strands. Data was fitted with a mono-exponential decay function. Binding half times match the fast species determined in (A). (C) Calculated off rates for different imager strands. Data points represent individual cells. Bars represent mean values for each condition and whiskers the standard deviation. $N_{replicates}$ = 1.*

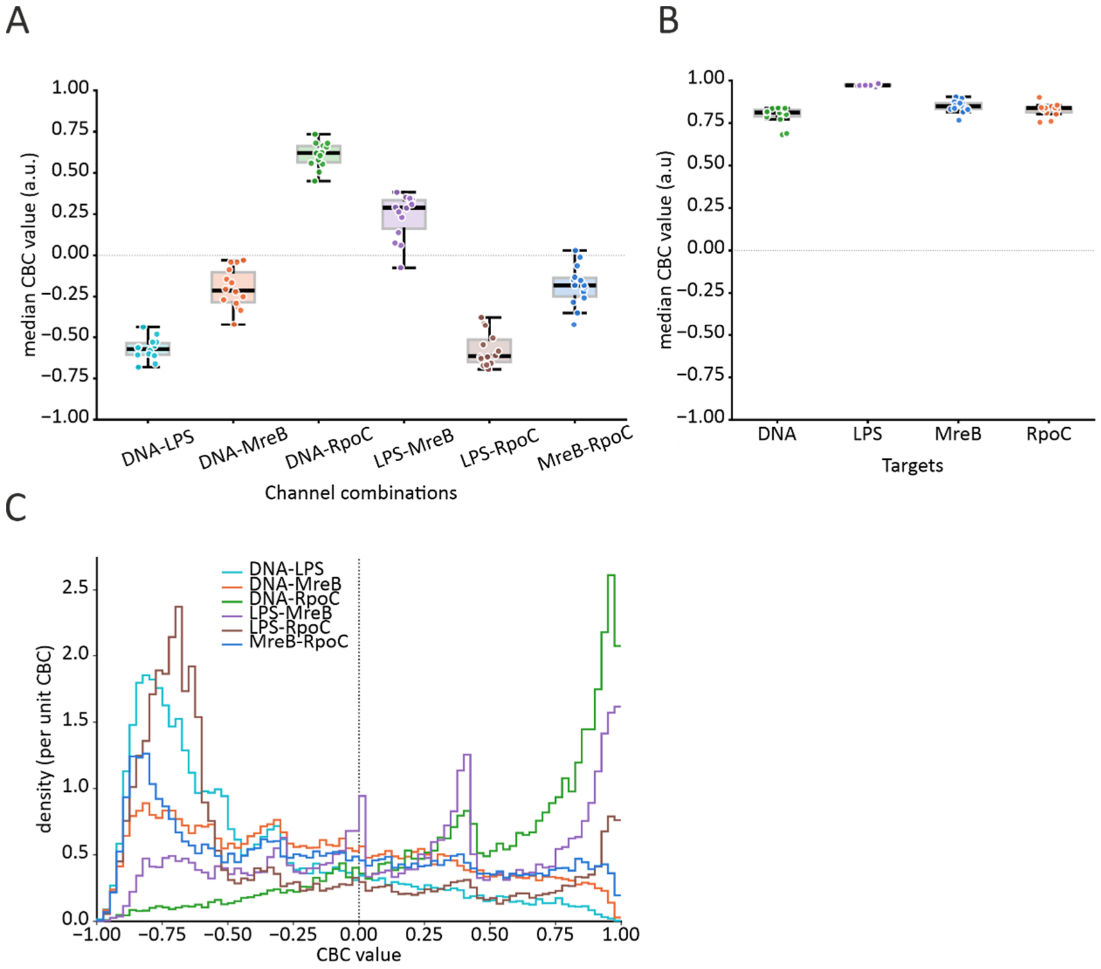


***Figure S8: Coordinate-based colocalization (CBC) of different channels of Exchange-PAINT measurements in E. coli****. (A) Median CBC values for different channel combinations. (B) Median CBC values for target positive controls. Localization lists were separated into two datasets (even and odd frame numbers) and subjected to CBC analysis. (C) CBC probability density distribution of the channel combinations shown in (A). Single-cell CBC data was pooled. $N_{cells}$ = 14, $N_{replicates}$ = 1.*

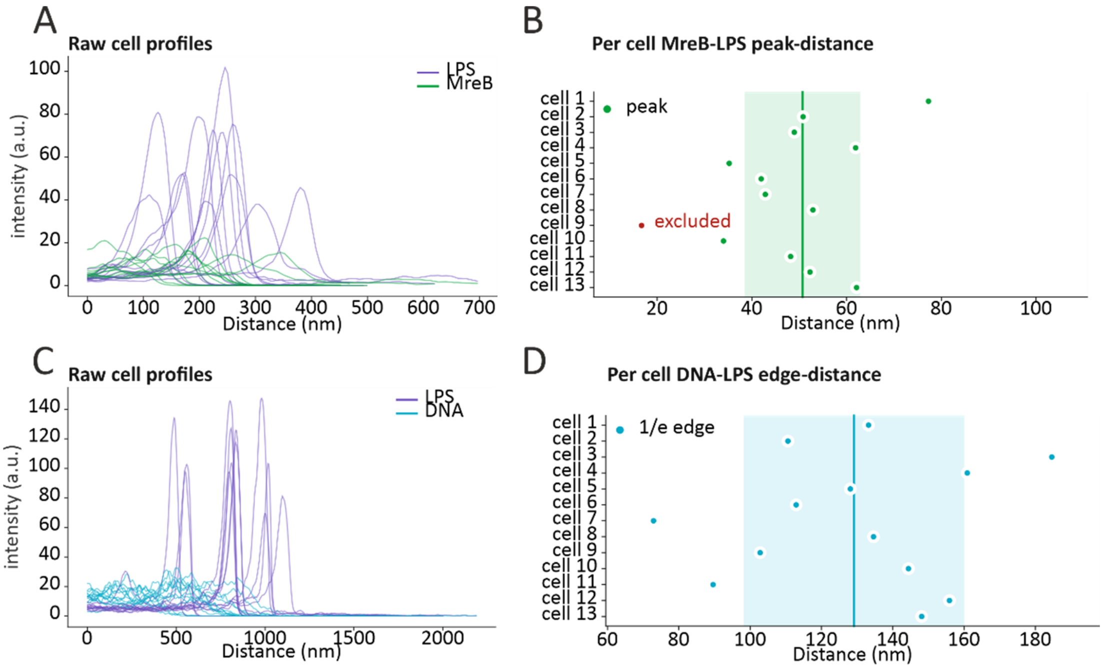


***Figure S9: Distance measurements in Exchange-PAINT measurements of E. coli.*** *(A) Raw profiles for LPS and MreB distributions determined for individual cells. Data was measured from cell cytosol towards the cell outside. (B) Peak-to-peak distances of LPS and MreB signals for individual cells. Cell 9 was excluded due to bad peak signal. The solid vertical line represents the mean value and shaded area the standard deviation. (C) Raw profiles of LPS and DNA distributions determined for individual cells. (D) Peak-to-1/e distance for LPS-DNA signals for individual cells. The solid vertical line represents the mean value and shaded area the standard deviation.* $N_{cells} = 13$.

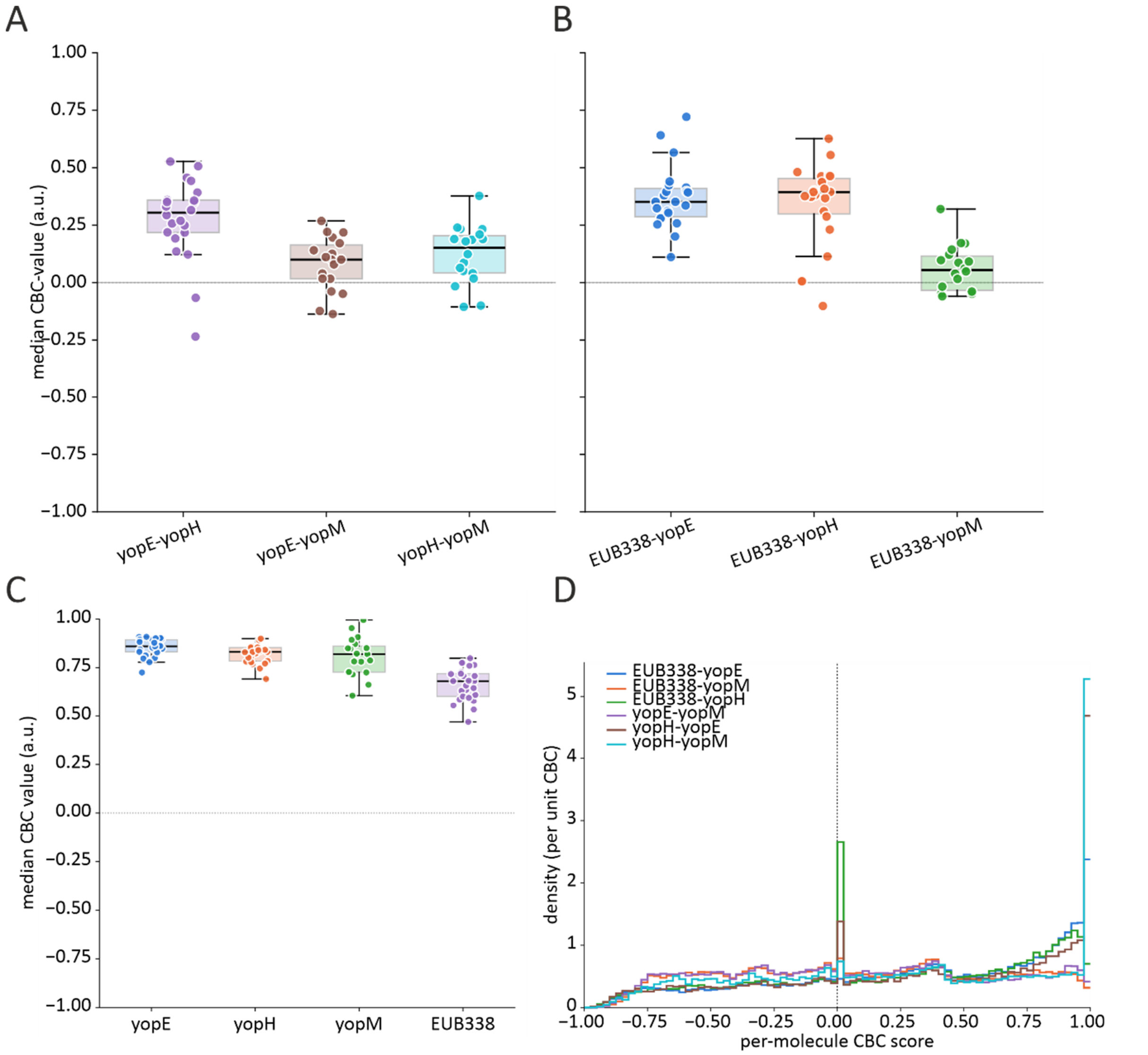


***Figure S10: Coordinate-based colocalization (CBC) analysis for Exchange-PAINT measurements in Y. enterocolitica***. *(A) CBC values for different effector mRNA combinations. (B) CBC values for different effector mRNAs against the EUB338 channel. (C) Median CBC values for target positive controls. Localization lists were separated into two datasets (even and odd frame numbers) and subjected to CBC analysis.* $N_{cells} = 13$.

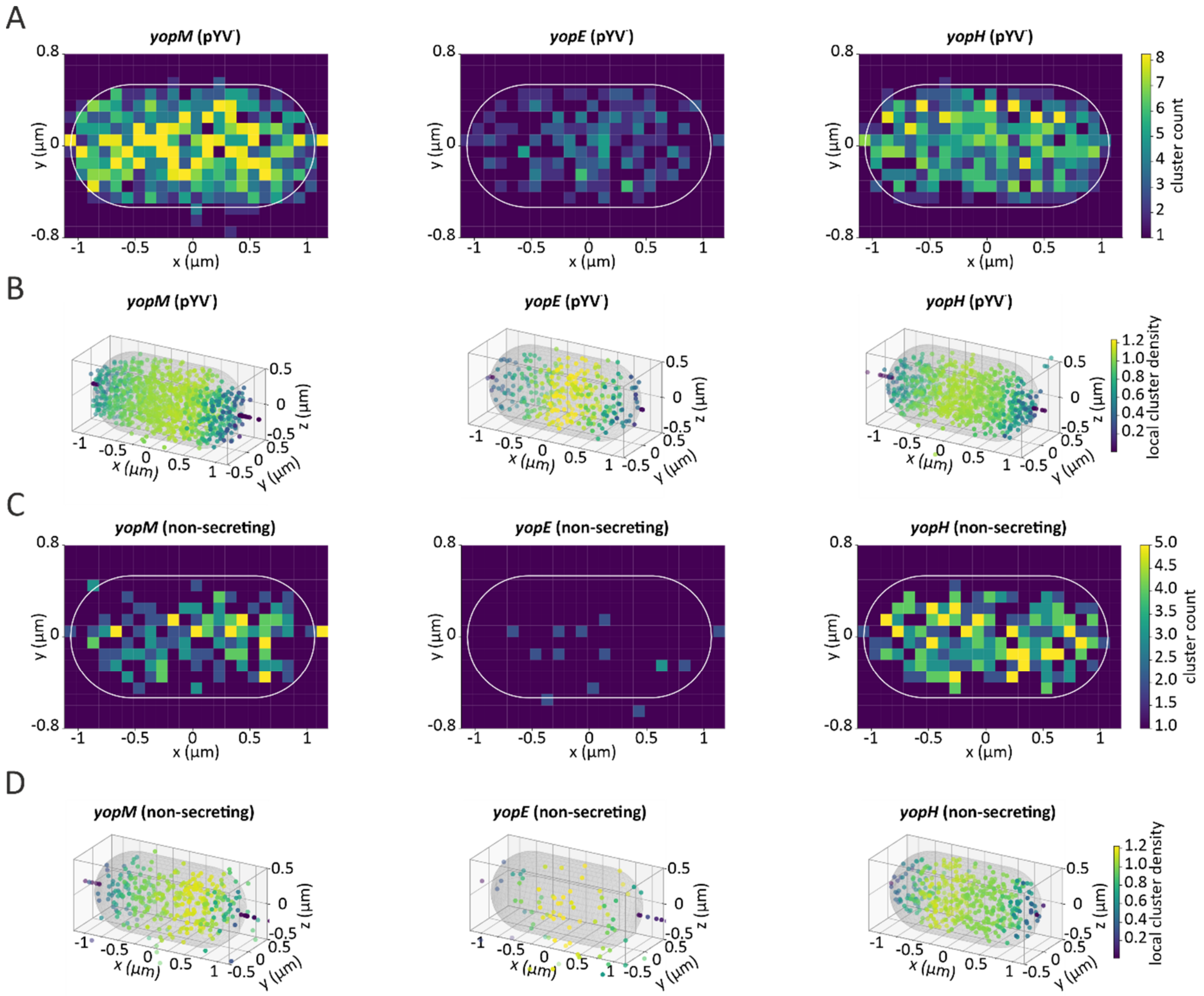


***Figure S 11: Cluster analysis of different effector mRNAs in Exchange-PAINT measurements in Y. enterocolitica****. (A) Normalized heatmaps of 2D-projected cluster distributions for different effector mRNAs in a Y. enterocolitica pYV ⁻ strain. Images are rendered by cluster count. (B) 3D distribution of the data shown in (A). Clusters are color-coded by local cluster density. (C) Normalized heatmaps of 2D-projected cluster distributions for different effector mRNAs in a Y. enterocolitica WT strain grown under non-secretion conditions. Images are rendered by cluster count. (D) 3D distribution of the data shown in (C). Clusters are color-coded by local cluster density. $N_{cells}$ = 58 (pYV ⁻) and 22 (WT, non-secreting).*

### Supplementary Tables

*Table S1: Bacterial strains and plasmids used in this study.*

| Strain | Genotype | Reference |
|---|---|---|
| MG1655 | *E. coli K-12* F⁻ λ⁻ *ilvG⁻ rfb-50 rph-1* | [2] |
| NO34 | MG1655, *mreB$^{sw}$-sfGFP csrD::kan* | [3] |
| ST18 | S17-1 *λpir ΔhemA* | [4] |
| pEB17 | Suicide plasmid for two-step allelic exchange, Kan$^{R}$, R6K-origin | [5] |
| pMBP-36 | pEB17 + *rpoC-ALFA* + homology arms | This study |
| pMBP-58 | pEB17 + *mreB-sfGFP-mreBCD* | This study |
| MBP-94 | *E. coli* MG1655 *rpoC-ALFA* | This study |
| MBP-128 | *E. coli* MG1655 *mreB$^{sw}$-sfGFP rpoC-ALFA* | This study |
| MRS40 | Wild-type *Y. enterocolitica E40, NalR, ΔblaA* | [6] |
| pYV$^{-}$ | MRS40 cured of virulence plasmid | [7] |

*Table S2: Summary of Oligos used in this study. DNA-PAINT docking sequences are shown in bold.*

| Oligo Name | Sequence | 5'-Mod | Used for |
|---|---|---|---|
| KA-3 | ATCGATCCTCTAGAGTCGACCTGCAGTAGAACGTGGTGACGTAATTTCCG | - | upstream homology arm for RpoC-ALFA |
| KA-17 | GTCAGACGACGACGCAGTTCTTCTTCCAGACGAGATATTTCTAGCTCGTTATCAGAACCGC | - | upstream homology arm for RpoC-ALFA |
| KA-8 | TGGAATTCCCGGGAGAGCTCAGATCCAAAGTAAAATGCCTTATCTCCTCC | - | downstream homology arm for RpoC-ALFA |
| KA-18 | AAATATCTCGTCTGGAAGAAGAACTGCGTCGTCGTCTGACCGAATAATCGTTAATCCGCAAATAACGTA | - | downstream homology arm for RpoC-ALFA |
| KA-21 | GATCTGAGCTCTCCCGG | - | Linearisation of pEB17 |
| KA-22 | TGCAGGTCGACTCTAGAG | - | Linearisation of pEB17 |
| KA-49 | CACCATCCGGATGTCCAAAAGGT | - | Validation Primer |
| KA-64 | GGAAGAAGAACTGCGTCGTC | - | Validation Primer |
| RX_36 | ATCGATCCTCTAGAGTCGACCTGCAATGTTGAAAAAATTTCGTGGCATG | - | MreB-sfGFP-MreBCD |
| RX_37 | TGGAATTCCCGGGAGAGCTCAGATCTTATTGCACTGCAAACTGC | - | MreB-sfGFP-MreBCD |
| RX_47 | GTGGTTGGTAAAGTAAGCG | - | Validation Primer |
| RX_48 | GTTGCGCAAGTAACTCC | - | Validation Primer |
| YE_yopH_R4x3_1 | ACCAATCGAGATACCTGACG**ACACACACACA** | - | yopH FISH-probe |
| YE_yopH_R4x3_2 | CTCTTAATTTCCCGGTACAA**ACACACACACA** | - | yopH FISH-probe |
| YE_yopH_R4x3_3 | TTCTTTATTGGCAGCAACGT**ACACACACACA** | - | yopH FISH-probe |
| YE_yopH_R4x3_4 | GGCTATGGTCAAACCTTGAA**ACACACACACA** | - | yopH FISH-probe |
| YE_yopH_R4x3_5 | ACGTGGCTTAGTACAGTTTG**ACACACACACA** | - | yopH FISH-probe |
| YE_yopH_R4x3_6 | GCGGTATCTTCTTGAGTTAG**ACACACACACA** | - | yopH FISH-probe |
| YE_yopH_R4x3_7 | AATTATGCTTTACCGTGCTT**ACACACACACA** | - | yopH FISH-probe |
| YE_yopH_R4x3_8 | TGCCGACACTTCTTAATTCA**ACACACACACA** | - | yopH FISH-probe |
| YE_yopH_R4x3_9 | AGTGTCATTTGGTCACTACG**ACACACACACA** | - | yopH FISH-probe |
| YE_yopH_R4x3_10 | GGTGCGTGAAGGACTGAATG**ACACACACACA** | - | yopH FISH-probe |
| YE_yopH_R4x3_11 | TCTGGGGTCTAGATGTGAAC**ACACACACACA** | - | yopH FISH-probe |

| | | | |
|---|---|---|---|
| YE_yopH_R4x3_12 | GGGCCATAAGGAGAAACAGT**ACACACACACA** | - | yopH FISH-probe |
| YE_yopH_R4x3_13 | GCTTGTAAGTAACGCGGATC**ACACACACACA** | - | yopH FISH-probe |
| YE_yopH_R4x3_14 | AATCGGTTTAGCTTTTCACC**ACACACACACA** | - | yopH FISH-probe |
| YE_yopH_R4x3_15 | TGCCGACAGCATTGAATATC**ACACACACACA** | - | yopH FISH-probe |
| YE_yopH_R4x3_16 | ACCTGGATGTAATTGGCATT**ACACACACACA** | - | yopH FISH-probe |
| YE_yopH_R4x3_17 | ACGCTATGGTACGAGTGTTA**ACACACACACA** | - | yopH FISH-probe |
| YE_yopH_R4x3_18 | GCATACGGAAATGGCTTTCA**ACACACACACA** | - | yopH FISH-probe |
| YE_yopH_R4x3_19 | TGGCTATCTCAGAACTGGAC**ACACACACACA** | - | yopH FISH-probe |
| YE_yopH_R4x3_20 | ATCTGGCATACCGAATCTTT**ACACACACACA** | - | yopH FISH-probe |
| YE_yopH_R4x3_21 | AGTGATACTGCCATAGGTAC**ACACACACACA** | - | yopH FISH-probe |
| YE_yopH_R4x3_22 | CCAACTTGCTGAGTCATTTT**ACACACACACA** | - | yopH FISH-probe |
| YE_yopH_R4x3_23 | AAATTGTTTTTTGACCCGCT**ACACACACACA** | - | yopH FISH-probe |
| YE_yopH_R4x3_24 | TAACTTCAGAGCTGACTGCG**ACACACACACA** | - | yopH FISH-probe |
| YE_yopH_R4x3_25 | GTTTTGTTTCTGCTGTTTGA**ACACACACACA** | - | yopH FISH-probe |
| YE_yopH_R4x3_26 | CTTCCTTTGCTTTCATACAT**ACACACACACA** | - | yopH FISH-probe |
| YE_yopH_R4x3_27 | AATGTATTACCGGCCGTAAT**ACACACACACA** | - | yopH FISH-probe |
| YE_yopH_R4x3_28 | ATTTGGCTGACCATATCTTC**ACACACACACA** | - | yopH FISH-probe |
| YE_yopH_R4x3_29 | GCTCATCTTTTTGTACCATA**ACACACACACA** | - | yopH FISH-probe |
| YE_yopH_R4x3_30 | CCTTCAGCCAACTTAATCAG**ACACACACACA** | - | yopH FISH-probe |
| YE_yopM_R3x3_1 | CTCAACTGGCATCTCAGTTA**CTCTCTCTCTC** | - | yopM FISH-probe |
| YE_yopM_R3x3_2 | CATTCCGACCATGCATTATA**CTCTCTCTCTC** | - | yopM FISH-probe |
| YE_yopM_R3x3_3 | ATCGCGTAACCTTGAAACCG**CTCTCTCTCTC** | - | yopM FISH-probe |
| YE_yopM_R3x3_4 | CCCCAGATTATTTAGTTCTA**CTCTCTCTCTC** | - | yopM FISH-probe |
| YE_yopM_R3x3_5 | AAATGCGGAGGTAATTCCGG**CTCTCTCTCTC** | - | yopM FISH-probe |
| YE_yopM_R3x3_6 | ACATGACGCCACTAAACTCT**CTCTCTCTCTC** | - | yopM FISH-probe |
| YE_yopM_R3x3_7 | GCAATTCCGGTAATTCTGTA**CTCTCTCTCTC** | - | yopM FISH-probe |
| YE_yopM_R3x3_8 | TCGGATAATGCCTTCAGATT**CTCTCTCTCTC** | - | yopM FISH-probe |
| YE_yopM_R3x3_9 | GGCACCTAAATATTCCAGTA**CTCTCTCTCTC** | - | yopM FISH-probe |
| YE_yopM_R3x3_10 | AGGACGAGTTTTGCAACTCT**CTCTCTCTCTC** | - | yopM FISH-probe |
| YE_yopM_R3x3_11 | CCAGCAGCAAGAAATTCCAG**CTCTCTCTCTC** | - | yopM FISH-probe |
| YE_yopM_R3x3_12 | GCAACTCTGACAATTCTTCC**CTCTCTCTCTC** | - | yopM FISH-probe |
| YE_yopM_R3x3_13 | ATCGCAGTCAAGAAGGGCAA**CTCTCTCTCTC** | - | yopM FISH-probe |
| YE_yopM_R3x3_14 | AAGTGTTTTCAGGGAAGGGG**CTCTCTCTCTC** | - | yopM FISH-probe |
| YE_yopM_R3x3_15 | CGGTAATTCTGGCAGATCAG**CTCTCTCTCTC** | - | yopM FISH-probe |
| YE_yopM_R3x3_16 | TGGCAATTCCGATAATCCAG**CTCTCTCTCTC** | - | yopM FISH-probe |
| YE_yopM_R3x3_17 | TTCATTGCTGGATGCATTGA**CTCTCTCTCTC** | - | yopM FISH-probe |
| YE_yopM_R3x3_18 | GTGAAGGGGGTAAATCGCAT**CTCTCTCTCTC** | - | yopM FISH-probe |
| YE_yopM_R3x3_19 | GGCAGTTCGATCAACTGATT**CTCTCTCTCTC** | - | yopM FISH-probe |
| YE_yopM_R3x3_20 | GTTCTAAGCGTGGAGGTAAC**CTCTCTCTCTC** | - | yopM FISH-probe |
| YE_yopE_R1x3_1 | AGGGGCAGTGATGTAGAAAT**TCCTCCTCCTCCT** | - | yopE FISH-probe |
| YE_yopE_R1x3_2 | CTAGATCCTGACACTGATGC**TCCTCCTCCTCCT** | - | yopE FISH-probe |
| YE_yopE_R1x3_3 | CAGACATTTCTCCTACGCTG**TCCTCCTCCTCCT** | - | yopE FISH-probe |

| | | | |
|---|---|---|---|
| YE_yopE_R1x3_4 | GCCAGATTGTTTGCATATTG**TCCTCCTCCTCCT** | - | yopE FISH-probe |
| YE_yopE_R1x3_5 | AATGATACGGCTGGCTAAGC**TCCTCCTCCTCCT** | - | yopE FISH-probe |
| YE_yopE_R1x3_6 | GAGTGGGCCATTGATGATAA**TCCTCCTCCTCCT** | - | yopE FISH-probe |
| YE_yopE_R1x3_7 | CCGAGAACATGCGTTGGATA**TCCTCCTCCTCCT** | - | yopE FISH-probe |
| YE_yopE_R1x3_8 | CATTTGTGCAGGCGTGAGTG**TCCTCCTCCTCCT** | - | yopE FISH-probe |
| YE_yopE_R1x3_9 | CACTGAAAGACGTAGGGCTT**TCCTCCTCCTCCT** | - | yopE FISH-probe |
| YE_yopE_R1x3_10 | GCAGCAAGTTGCTTGATACT**TCCTCCTCCTCCT** | - | yopE FISH-probe |
| YE_yopE_R1x3_11 | CATGTATTTTGGCAGCGTCT**TCCTCCTCCTCCT** | - | yopE FISH-probe |
| YE_yopE_R1x3_12 | CTGCATCCAAGCTACTCAAC**TCCTCCTCCTCCT** | - | yopE FISH-probe |
| YE_yopE_R1x3_13 | TGGCGAACTGGTCATGATTT**TCCTCCTCCTCCT** | - | yopE FISH-probe |
| YE_yopE_R1x3_14 | TTGGCATTGAGTGATACTGC**TCCTCCTCCTCCT** | - | yopE FISH-probe |
| YE_yopE_R1x3_15 | CCACCACAAAACTGCATCAG**TCCTCCTCCTCCT** | - | yopE FISH-probe |
| YE_yopE_R1x3_16 | TTTAAAATGGCACTGGCCTC**TCCTCCTCCTCCT** | - | yopE FISH-probe |
| YE_yopE_R1x3_17 | GGGAATACCACAAACAGGCG**TCCTCCTCCTCCT** | - | yopE FISH-probe |
| YE_yopE_R1x3_18 | CGTTAGATCAACGCCACTGG**TCCTCCTCCTCCT** | - | yopE FISH-probe |
| YE_yopE_R1x3_19 | CAGCCCTTTGATCTCATTTG**TCCTCCTCCTCCT** | - | yopE FISH-probe |
| YE_yopE_R1x3_20 | CAGCCCTTTGATCTCATTTG**TCCTCCTCCTCCT** | - | yopE FISH-probe |
| Cy3-EUB338-R1x3 | GCTGCCTCCCGTAGGAGT**TCCTCCTCCTCCT** | Cy3 | 16S subunit rRNA FISH-probe |
| Cy3-EUB338-R2x3 | GCTGCCTCCCGTAGGAGT**ACCACCACCACCA** | Cy3 | 16S subunit rRNA FISH-probe |
| Cy3-EUB338-R3x3 | GCTGCCTCCCGTAGGAGT**CTCTCTCTCTC** | Cy3 | 16S subunit rRNA FISH-probe |
| Cy3-EUB338-R4x3 | GCTGCCTCCCGTAGGAGT**ACACACACACA** | Cy3 | 16S subunit rRNA FISH-probe |
| Cy3-EUB338-R5x3 | GCTGCCTCCCGTAGGAGT**CTTCTTCTTCTTC** | Cy3 | 16S subunit rRNA FISH-probe |
| Cy3-EUB338-R6x3 | GCTGCCTCCCGTAGGAGT**AACAACAACAACAA** | Cy3 | 16S subunit rRNA FISH-probe |
| Cy3-EUB338-R1x3_scrambled | CCGGCATTTGGATGCGCC**TCCTCCTCCTCCT** | Cy3 | 16S subunit rRNA FISH-probe |
| Cy3-EUB338-R2x3_scrambled | CCGGCATTTGGATGCGCC**ACCACCACCACCA** | Cy3 | 16S subunit rRNA FISH-probe |
| Cy3-EUB338-R3x3_scrambled | CCGGCATTTGGATGCGCC**CTCTCTCTCTC** | Cy3 | 16S subunit rRNA FISH-probe |
| Cy3-EUB338-R4x3_scrambled | CCGGCATTTGGATGCGCC**ACACACACACA** | Cy3 | 16S subunit rRNA FISH-probe |
| Cy3-EUB338-R5x3_scrambled | CCGGCATTTGGATGCGCC**CTTCTTCTTCTTC** | Cy3 | 16S subunit rRNA FISH-probe |
| Cy3-EUB338-R6x3_scrambled | CCGGCATTTGGATGCGCC**AACAACAACAACAA** | Cy3 | 16S subunit rRNA FISH-probe |
| Cy3-EUB338-R7x3 | GCTGCCTCCCGTAGGAGT **TTCCTTCCTTCCTTCC** | Cy3 | 16S subunit rRNA FISH-probe |
| Cy3-EUB338-R8x3 | GCTGCCTCCCGTAGGAGT**AACCAACCAACCAACC** | Cy3 | 16S subunit rRNA FISH-probe |
| Cy3-EUB338-R9x3 | GCTGCCTCCCGTAGGAGT**CCTCCCTCCCTCCCTC** | Cy3 | 16S subunit rRNA FISH-probe |

| | | | |
|---|---|---|---|
| Cy3-EUB338-R10x3 | GCTGCCTCCCGTAGGAGT**CCCACCCACCCACCCA** | Cy3 | 16S subunit rRNA FISH-probe |
| Cy3-EUB338-R11x3 | GCTGCCTCCCGTAGGAGT**TTTCTTTCTTTCTTTC** | Cy3 | 16S subunit rRNA FISH-probe |
| Cy3-EUB338-R12x3 | GCTGCCTCCCGTAGGAGT**AACAAACAAACAAACA** | Cy3 | 16S subunit rRNA FISH-probe |
| Cy3-EUB338-R7x3-scrambled | CCGGCATTTGGATGCGCC**TTCCTTCCTTCCTTCC** | Cy3 | 16S subunit rRNA FISH-probe |
| Cy3-EUB338-R8x3-scrambled | CCGGCATTTGGATGCGCC**AACCAACCAACCAACC** | Cy3 | 16S subunit rRNA FISH-probe |
| Cy3-EUB338-R9x3-scrambled | CCGGCATTTGGATGCGCC**CCTCCCTCCCTCCCTC** | Cy3 | 16S subunit rRNA FISH-probe |
| Cy3-EUB338-R10x3-scrambled | CCGGCATTTGGATGCGCC**CCCACCCACCCACCCA** | Cy3 | 16S subunit rRNA FISH-probe |
| Cy3-EUB338-R11x3-scrambled | CCGGCATTTGGATGCGCC**TTTCTTTCTTTCTTTC** | Cy3 | 16S subunit rRNA FISH-probe |
| Cy3-EUB338-R12x3-scrambled | CCGGCATTTGGATGCGCC**AACAAACAAACAAACA** | Cy3 | 16S subunit rRNA FISH-probe |
| Cy5-EUB338 | GCTGCCTCCCGTAGGAGT | Cy5 | 16S subunit rRNA FISH-probe |
